# High dimensional flow cytometric analysis reveals distinct NK cell subsets but conserved response to cytokine stimulation in umbilical cord blood and adult peripheral blood

**DOI:** 10.1101/2022.07.19.500730

**Authors:** Irina Buckle, Vicky Weinert, David P. Sester, Kristen Radford, Camille Guillerey

**Affiliations:** Cancer Immunotherapies Laboratory, Mater Research Institute, University of Queensland, Translational Research Institute, Brisbane, Queensland, Australia; TRI Flow cytometry suite, Translational Research Institute, 37 Kent Street, 4102 Woolloongabba QLD, Australia

## Abstract

Adoptive cellular therapies using Natural Killer (NK) cells are of growing interest. NK cells can be obtained from various sources, including umbilical cord blood (UCB) and adult peripheral blood (APB). Understanding the diversity of NK cell populations and their receptor expression in both UCB and APB will guide future therapeutic designs. In this study, we used a 20-colour flow cytometry panel to compare unstimulated and cytokine-activated UCB and APB NK cells. Our analysis showed that UCB NK cells express slightly higher levels of the immune checkpoints PD-1, TIGIT and CD96 compared to their APB counterparts. Unsupervised hierarchical clustering and principal component analyses revealed previously unappreciated differences in NK cell populations from UCB and APB. UCB was characterised by an enrichment in CD56^neg^ as well as mature NKp46^neg^ and CD56^+^CD16^+^ NK cell populations whereas CD57^+^ terminally differentiated NK cells with variable expression of KIRs and CD16 were found in APB. These populations were conserved following two-days of IL-15 culture as well as overnight stimulation with IL-12, IL-15, and IL-18. Interestingly, cytokine stimulation was associated with the up-regulation of LAG-3 and DNAM-1 together with the downregulation of NKG2D, TIGIT and CD16 on multiple NK cell subsets in both UCB and APB. TIM-3 was also up-regulated with activation, but only in UCB. Overall, our data indicate that NK cells in UCB have a more immature phenotype than APB NK cells and UCB NK cells might be more amenable to immune checkpoint therapy.

## Introduction

Natural Killer (NK) cells are innate lymphoid cells (ILCs) controlling cancer and infections (1). NK cells utilise an array of germline-encoded receptors to detect potentially dangerous cells such as damaged, malignant or virally infected cells (2). NK cells release cytotoxic granules containing granzymes and perforin resulting in the lysis of target cells. NK cells also exert their protection through the secretion of cytokines, mainly IFN-γ and TNF-α as well as a broad range of pro-inflammatory chemokines that contribute to the shaping of innate and adaptive immune responses (3).

Human peripheral NK cells are commonly divided into two populations. CD56^dim^ NK cells express the Fc gamma receptor III (CD16) and have been described to be highly reactive to target cells with high cytotoxic capabilities (4). By contrast, CD56^bright^ NK cells are CD16^low^. CD56^bright^ NK cells are responsive to cytokine stimulation and are strong producers of IFN-γ. Moreover, when primed with IL-15, CD56^bright^ NK cells show potent response to tumour cells (5). According to the linear model of NK cell differentiation, CD56^bright^ NK cells expressing high levels of the inhibitory receptor NKG2A and low levels of KIR receptors differentiate into CD56^dim^ NK cells acquiring KIR expression and reduced levels of NKG2A (6). The acquisition of CD57 on CD56^dim^ NK cells marks their terminal differentiation (7). Moreover, we recently discovered an alternative pathway of terminal maturation of human adult peripheral blood (APB) NK cells characterised by the loss of DNAM-1 (CD226) expression (8). Overall, it appears evident that diverse NK cell subsets with distinct phenotypic characteristic also harbors specific functional properties which differently contribute to the maintenance of the body homeostasis. However, the scope of this diversity is still not fully understood in humans (9).

Allogeneic NK cell therapies have proved safe and shown promising results in cancer patients (10, 11). Sources of allogeneic NK cells include APB, umbilical cord blood (UCB), stem cells or NK cell lines (12). UCB is a particularly attractive source considering the ease of access, cryopreservation possibility and the reported high numbers of UCB NK cells (13–15). Previous studies have reported differences between APB and UCB, with UCB NK cells expressing reduced levels of L-selectin, CD57, KIRs, DNAM-1 and NKG2C but higher levels of NKG2A (16, 17). However, these studies have been limited by the low number of markers examined by flow cytometry analysis, not allowing for co-expression assessment and a complete delineation of NK cell subsets.

The ability of immune checkpoints to regulate NK cell functions has recently attracted great interest as these molecules could be clinically targeted to further enhance NK cell functions (18). Previous studies have reported expression of TIGIT, CD96, TIM-3, LAG-3 and PD-1 (8, 19–22) while data reporting CTLA-4 expression remain controversial (23). To date, immune checkpoints on UCB NK cells remain uncharacterised.

In this study, we employed a 20-colour flow cytometry panel to compare immune checkpoint expression and NK cell subsets between UCB and APB, either unstimulated or upon culture with cytokines. Our data reveals previously overlooked NK cell subsets that differ between UCB and APB. Moreover, we demonstrate higher levels of immune checkpoint expression on UCB NK cells when compared to APB. Overall, our study highlights fundamental differences between UCB and APB NK cells that should be considered in the choice of NK cell source for adoptive cellular therapy.

## Material and Methods

### Samples

Adult peripheral blood (APB) samples were collected from healthy individuals following written consent in line with standards established by the Declaration of Helsinki. Umbilical cord blood (UBC) samples were provided by Queensland Cord Blood Bank. All human sample collection was approved by Mater Human Research Ethics Committee (HREC/13/MHS/83 and 22310/1407AP).

### Stimulation

Mononuclear cells were cultured for 36 hrs at 37°C, 5% CO2 with 50 ng/ml animal-free recombinant human IL-15 (Peprotech). For stimulation, 10ng/ml, human IL-12p70 (HEK293) (Peprotech) and 50ng/ml recombinant human IL-18 (MBL International) were added to the IL-15 cultures for the last 12 hrs (overnight stimulation).

### Flow Cytometry

Non-specific antibody binding was prevented by blocking cells with 10% rat and 10% mouse serum for 10 minutes at 4°C. Cells were washed in PBS and stained with fixable viability stain FVS700 (BD Biosciences) for 10 minutes at 4°C, protected from light. Cells were subsequently washed in autoMACS running buffer (Miltenyi Biotec Inc.) and centrifuged at 500g for 5 minutes at 4°C. Labelling with fluorescent antibodies was performed for 25 minutes, at room temperature, protected from light. Antibody mixes were prepared in brilliant stain buffer (BD Biosciences). The details of antibodies and dilutions used are included Supplementary Table 1. Samples were acquired on a BD FACSymphony A5.

### Analysis

Data was analysed using Flowjo v10.8.1 or v10.7.2. For hierarchical clustering, FlowSOM pluggin v2.2.0 was used together with R v4.1.2. Statistical analyses were performed using Graphpad Prism v8.0.1.

## Results

### UCB NK cells express high levels of CD96, TIGIT and TIM-3

We applied a 20-colour flow cytometry panel (Supplementary Table 1) to define the expression of immune checkpoints and NK cell receptors in UBC NK cells compared to APB NK cells. As per previous studies (24), NK cells were defined as CD3^-^CD33^-^CD19^-^ cells that expressed either CD56 or CD16 (Supplementary Figure 1A). NK cells represented 6.6 + 3.6 % of human CD45^+^ cells in UCB and 6.1 + 3.4 % in APB (Supplementary Figure 1B). In agreement with published reports (16, 17), we found that UCB NK cells contained a significantly higher frequency of NKG2A^+^ cells and a lower percentage of CD57^+^ cells compared to APB NK cells (Supplementary Figure 1C). We did not observe any significant difference in the frequencies of NKp46^+^, CD16^+^, KIR^+^ or NKp30^+^ NK cells between UCB and APB (Supplementary Figure 1C). By contrast, the frequency of NKG2D^+^ NK cells was increased in UCB while the frequency of DNAM-1^+^ (CD226^+^) NK cells was lower in UCB compared to APB (Figures 1A-B). We did not detect expression of the immune checkpoints CTLA-4 or LAG-3 on unstimulated NK cells (data not shown). TIM-3 was expressed on approximately 60% of NK cells in both APB and UCB, with similar expression levels (Figures 1C-D). Interestingly, CD96 and TIGIT were expressed in a higher proportion of UCB NK cells compared with APB NK cells (Figures 1C-D). Moreover, a small percentage of UCB NK cells expressed PD-1 (Figure 1E). Compared to their PD-1-negative counterpart, PD-1^+^ UCB NK cells were mostly CD16^+^ and expressed slightly increased levels of KIRs and NKp44, indicating a mature/activated phenotype (Figure 1F). However, UCB PD-1^+^ NK cells displayed reduced expression of other NK cell receptors including NKG2A, NKG2D, NKp30, DNAM-1 (Supplementary Figure 1D). Half of UCB PD-1^+^ NK cells were negative for CD56 (Figure 1F). PD-1^+^ UCB NK cells also expressed very low levels of the terminal differentiation marker CD57. Finally, UCB PD-1^+^ NK cells showed reduced expression of TIM-3 and TIGIT, indicating that these immune checkpoints are not co-expressed. These data collectively establish that UCB NK cells express higher levels of immune checkpoints than APB NK cells, with TIM-3, TIGIT and CD96 being expressed on a large proportion of unstimulated NK cells. In addition, PD-1 is expressed a higher frequency of UCB NK cells compared to APB counterparts.

**Figure 1.**
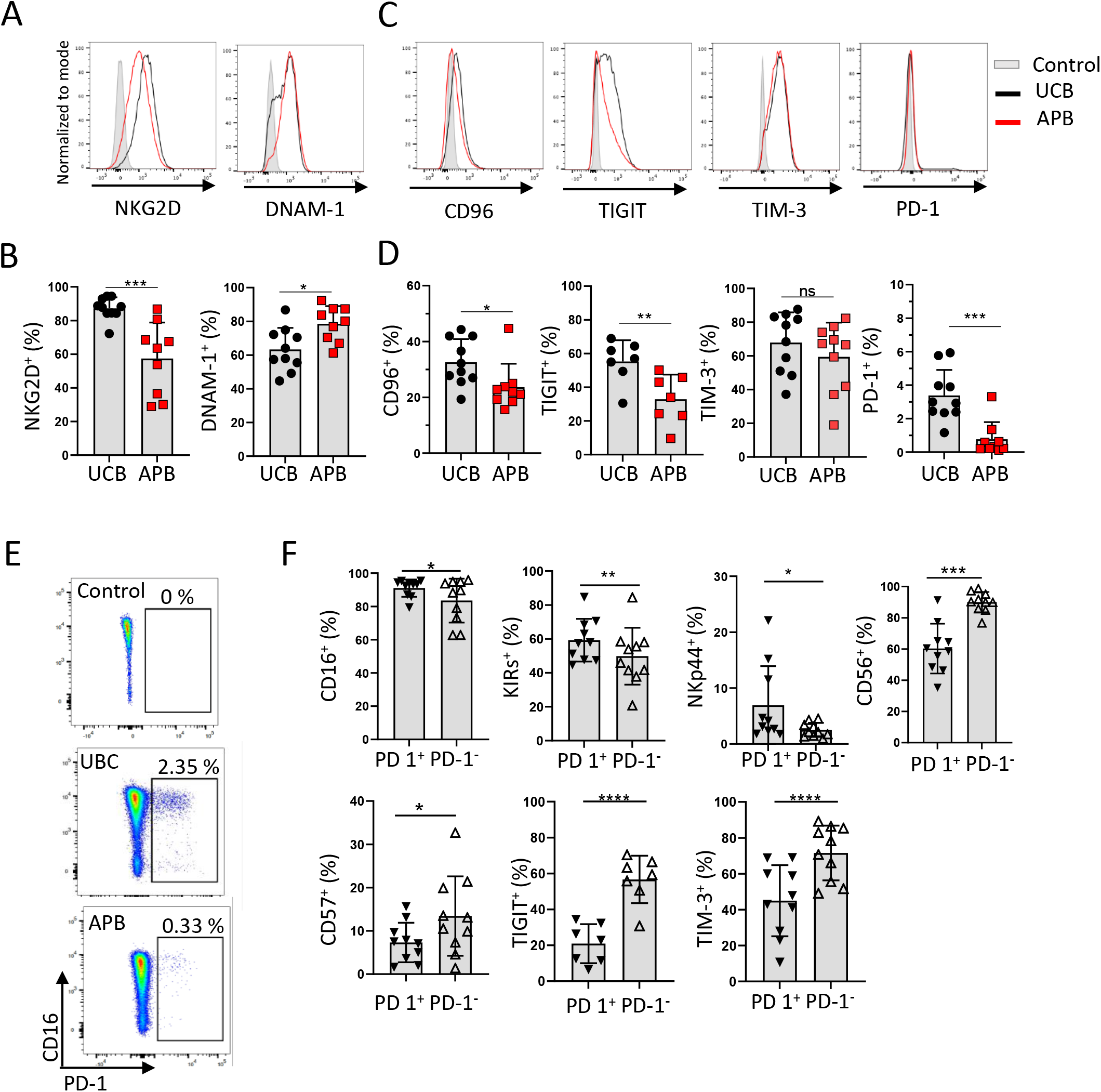
UCB NK cells express higher levels of immune checkpoints compared to APB NK cells. UCB MNCs were stained with a 20-colour flow cytometry panel and individual receptor expression were analysed on NK cells. **(A-C)** Representative histograms showing receptor expression on UCB NK cells (black) and APB NK cells (red). Filled histogram show UCB control sample stained with only NK cell defining markers **(B-D)** Graphs show percentages of NK cells positive for a given marker in UCB and APB. **(E)** Representative dot-plots showing PD-1 expression on NK cells. **(F)** Percentages of PD-1^+^ and PD-1^-^ UCB NK cells positive for a given marker. Data are from 3 independent experiments with a total n=10 UCB and n=9 APB samples. Graphs show mean + SD. Data were analysed with a paired (F) or unpaired (B-D) t-test. ns = non significant; * p<0.05; ** p<0.01; *** p<0.001; **** p <0.0001

### UCB NK cells contain an unconventional population of CD56-negative NK cells

To further characterise UCB NK cells, we divided NK cells into four subsets according to their expression of CD56 and CD16: CD56^bright^CD16^-^, CD56^+^CD16^+^, CD56^dim^CD16^-^ and CD56^-^ CD16^+^ (hereafter referred as CD56^neg^) NK cells (Figure 2A). When quantified in relation to live CD45^+^ cells, the three CD56^+^ populations (CD56^bright^CD16^-^, CD56^+^CD16^+^ and CD56^dim^CD16^-^ and NK cells) were found at similar frequencies in UCB and APB (Supplementary Figure 2A) while frequencies of CD56^neg^ NK cells were significantly increased in UCB (Figure 2B). Quantified as proportions of NK cells, CD56^neg^ NK cells represented 10.5 + 6.2 % of UCB NK cells but were virtually absent of APB NK cells (Figure 2C). Extended phenotypic analysis of CD56^neg^ NK cells revealed significantly decreased frequencies of cells positive for the activating receptors DNAM-1, NKG2D and NKp30 when compared to all UBC NK cells (Figure 2D). Interesting, CD56^neg^ NK contained fewer cells expressing both the terminal differentiation marker CD57 and the inhibitory receptor NKG2A which is classically expressed on immature CD56^bright^ NK cells. This might indicate an intermediate maturation status of CD56^neg^ NK cells. The frequencies of NKp46^+^, KIR^+^, TIM-3^+^ and CD96^+^ cells were all similar when comparing the CD56^neg^ UCB NK population to the whole UCB NK cells compartment (Supplementary Figure 2B). Finally, relatively to all UBC NK cells we observed a significantly reduced prevalence of TIGIT^+^ cells but a significantly higher expression frequency of PD-1^+^ cells in UCB CD56^neg^ NK cells (Figure 2D). We concluded that UCB contains an expanded population of CD56^neg^ NK cells expressing the NK cell markers CD16, NKp46 and KIRs and enriched in PD-1 ^+^ NK cells.

**Figure 2.**
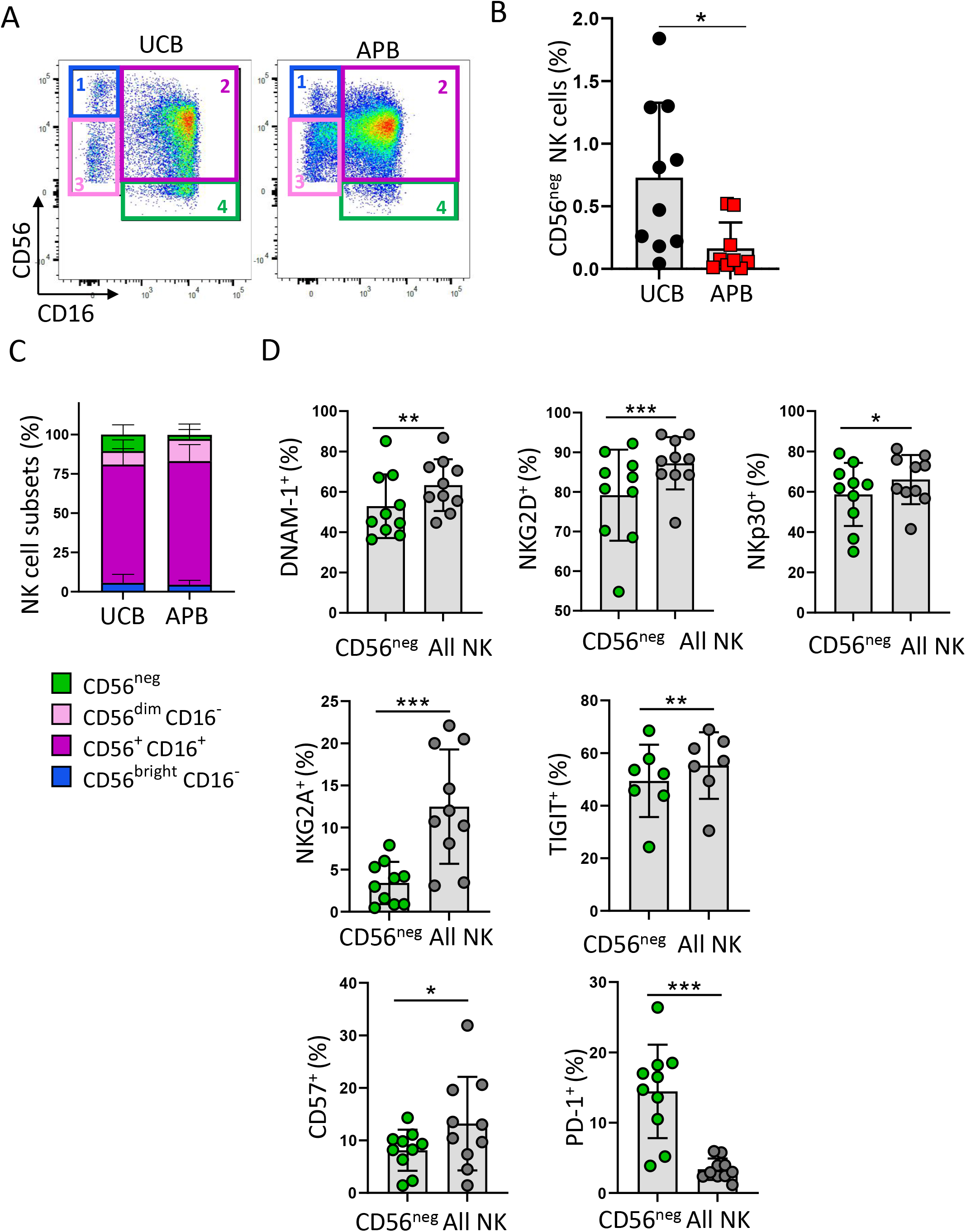
UCB contains a population of CD56^neg^ NK cells with increased PD-1 expression. **(A)** Representative gating strategy showing CD56^bright^CD16^-^ (1), CD56^+^CD16^+^ (2) CD56^dim^CD16^-^ (3) and CD56^neg^ (4) NK cell subsets in UCB and APB. **(B)** Graph showing the percentage of CD56^neg^ NK cells out of live human CD45^+^ cells in UCB and APB. **(C)** Graph displays the contribution of each identified NK cell subset within UCB and APB NK cell compartments. Data was analysed using a 2-way ANOVA (UCB/APB vs subset) followed by a Sidak’s multiple comparison test. CD56^neg^ subset was found differentially represented in UCB compared to APB (p=0.0144). No significant difference was observed for the 3 other subsets. **(D)** Percentages of UCB CD56^neg^ NK cells positive for a given marker compared to the whole UCB NK cell population. Data are from 3 independent experiments with a total n=10 UCB and n=9 APB samples. Graphs show mean + SD. Data were analysed with a paired (D) or unpaired (B-C) t-test. ns = non-significant; * p<0.05; ** p<0.01; *** p<0.001; **** p <0.0001

### High-dimensional analysis reveals additional unconventional NK cell subsets in UCB and APB

We applied the unsupervised hierarchical clustering algorithm FlowSOM (25) to our data. FlowSOM was run on a first set of three UCB and three APB samples, leading to the identification of 16 clusters (Supplementary Figure 3A). Clusters were then grouped into 8 main populations named A to H (Supplementary Figure 3B). Populations A-G were also identified in a second independent analysis performed on four UCB and four APB samples (Supplementary Figure 3C). FlowSOM minimum spanning tree highlighting the identified populations is shown in Figure 3A and populations were overlayed on a TriMAP principal component analysis (Figure 3B). Figure 3C shows TriMAP heatmaps of the markers used to delineate the populations. Population A corresponded to CD56^bright^ NK cells and was subdivided into A_1_ (CD56^bright^CD16^-^) which matched the conventional description of the CD56^bright^ population and A_2_ (CD56^bright^CD16^+^) which might constitute an intermediate population between CD56^bright^ and CD56^dim^ NK cells. Population B regrouped conventional mature NK cells (NKp46^+^CD56^dim^CD16^+^) with B_1_ corresponding to mature CD57^-^ NK cells and B_2_ corresponding to terminally differentiated CD57^+^ NK cells. Population C corresponded to CD56^neg^ NK cells and was further sub-divided into 3 subsets: C_1_, the most abundant CD56^neg^ subset was NKp46^+^KIR^-^; C_2_ was a small subset of NKp46^-^KIR^-^CD56^-^ NK cells that were positive for PD-1 and C_3_ regrouped NKp46^+^KIR^+^CD56^-^ NK cells. Population D was constituted of CD56^+^CD16^+^ NK cells that might be a transitioning subset between CD56^bright^CD16^+^ and CD56^dim^CD16^+^ as these cells were negative for KIRs. Population E corresponded to mature (CD16^+^KIR^+^) NK cells that were negative for NKp46, with E_1_ being PD-1^-^ and E_2_ being PD-1^+^. Population F encompassed unconventional CD56^dim^CD16^-^ NK cells with F1 being KIR^+^ and F2 being KIR^-^. Population G regrouped unconventional terminally differentiated (CD57^+^) NK cells with G_1_ being negative for CD16, G_2_ being negative for KIRs and G_3_ being negative for both CD16 and KIRs. Finally, population H corresponded to unconventional CD56^bright^ NK cells that were also positive for KIRs; however this subset was not identified in the second analysis on a different dataset (Supplementary Figure 3C). In summary, unsupervised hierarchical clustering enabled the identification of novel NK cells subsets such as unconventional terminally differentiated NK cells that have previously not been reported in the literature.

**Figure 3.**
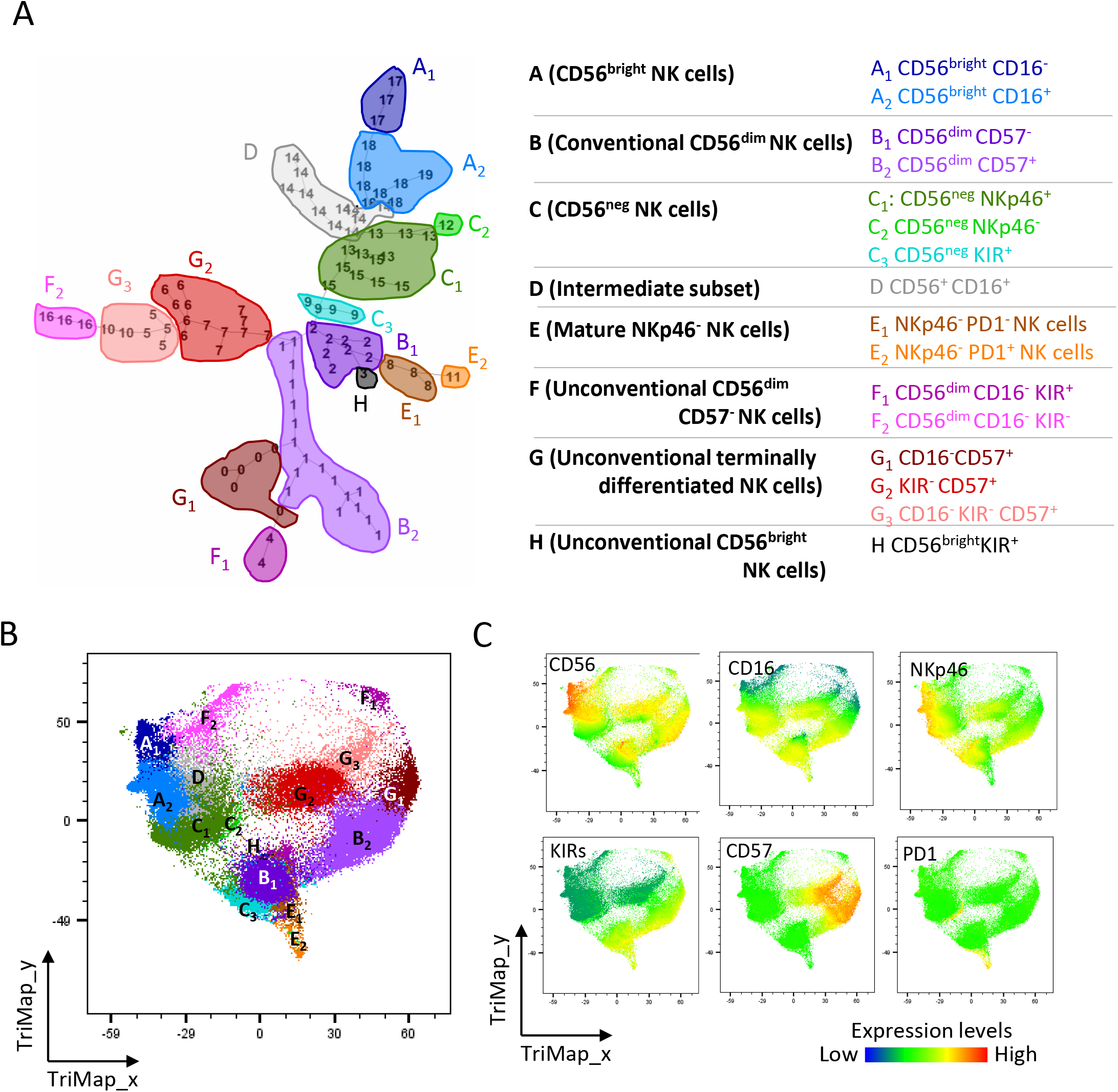
High dimensional analysis identifies unconventional NK cell populations. **(A)** FlowSOM algorithm was run on NK cells from 3 UCB and 3 APB samples. Populations as identified in Supplementary Figure 3A are shown on the Flowjo generated tree. **(B)** TriMap algorithm was run on the same dataset and FlowSOM populations were overlaid. **(C)** Heatmaps showing expression levels of NK cell receptors used to identify NK cells subsets.

### High-dimensional analysis highlights different representation of NK cell populations in UCB and APB

We then analysed the distribution of the newly identified subsets in UCB and APB. TriMAP analysis indicated that the terminally differentiated NK cell subsets G_1_, G_3_ and B_2_ consisted mainly of APB NK cells whilst CD16^+^KIR^+^NKp46^-^ NK cells (Population E) were mostly comprised of UCB NK cells (Figure 4A). Quantitative analysis of populations indicated that CD56^bright^ NK cells (Population A) were twice more abundant in UCB than in APB (15.4% and 8.4 % of all NK cells, respectively; Figure 4B). Conventional CD56^dim^ NK cells (Population B) represented more than one quarter of all NK cells in both UCB and APB. In line with the less mature profile of UCB NK cells (17), the CD57^-^ subset B1 was more represented in UCB while the terminally differentiated subset B2 was mostly found in APB (Figure 4B). Confirming data from Figure 2, CD56^neg^ NK cells (Population C) accounted for a higher percentage of NK cells in UCB, in comparison with APB. Intermediate CD56^+^CD16^+^ NK cells were also more represented in UCB compared to APB (17.9% and 8.8%, respectively). Unconventional mature NK cells that were either NKp46^-^ or CD16^-^ (Populations E and F) constituted rather small subsets (5% or less) of NK cells in both UCB and APB. Finally, unconventional terminally differentiated subsets represented more than a quarter of APB NK cells whereas they were poorly represented in UCB. CD56^bright^ KIR^+^ NK cells (Population H) constituted less than 1% of NK cells in both UCB and APB. An independent analysis on a second set of samples confirmed these results, the only difference being a higher representation of population F in the second data set (Supplementary Figure 4). Taken together, these data indicate that conventional NK cell subsets (e.g. CD56^bright^, Population A; and CD56^dim^, Population B) NK cells constitute less than half of all NK cell subsets in both UCB and APB. We identified unconventional NK cell subsets that were differently represented in UCB and APB; terminally differentiated NK cells that lack CD16 and/or KIR expression (Population G) were abundant in APB while CD56^neg^ NK cells (Population C), CD56^+^CD16^+^ NK cells (Population D) as well as mature NKp46^-^ NK cells (population E) were more represented in UCB.

**Figure 4.**
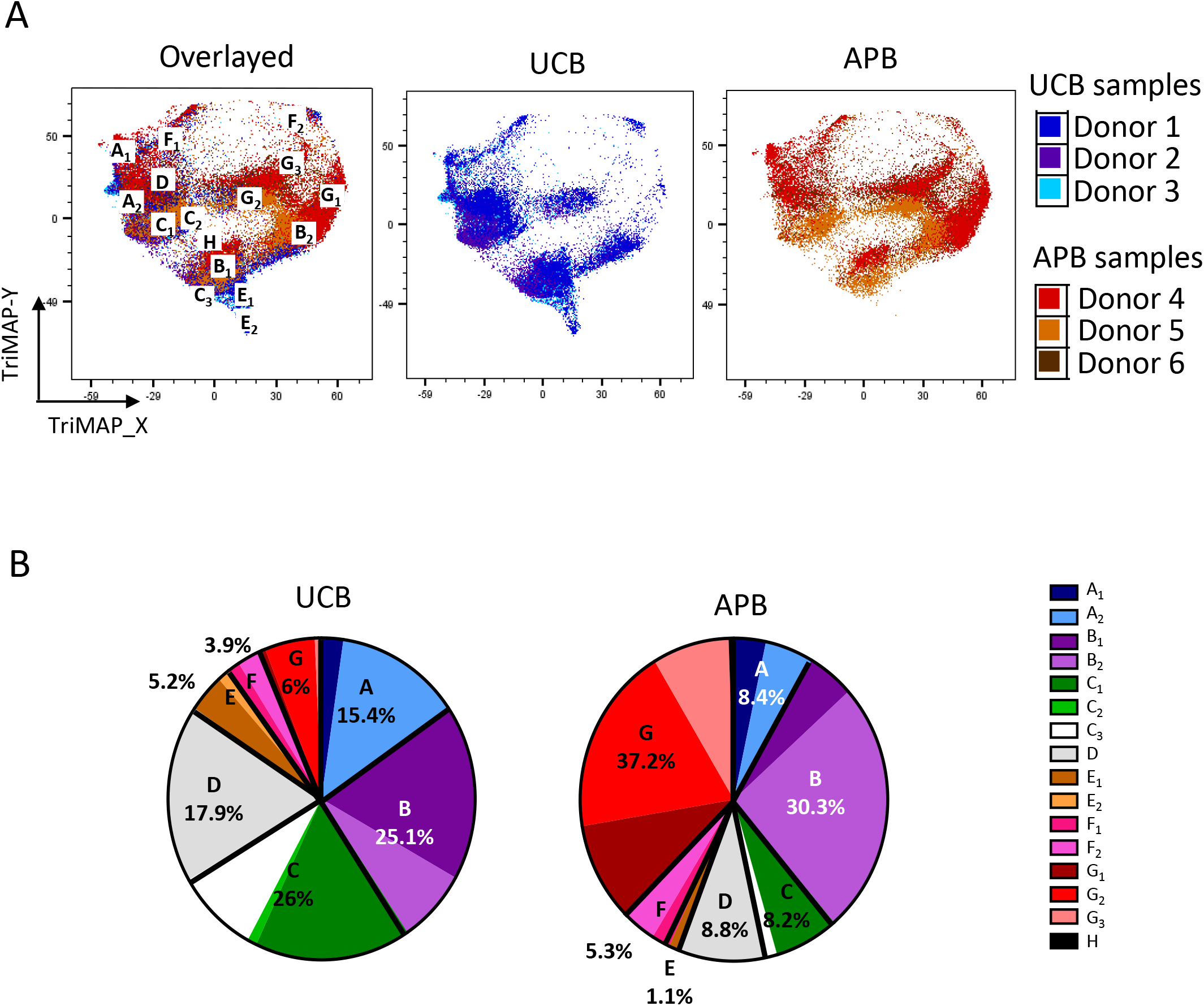
Different representation of NK cell populations in UCB and APB. **(A)** Individual donor contribution is shown on TriMap dot plot using the dataset analysed in Figure 3. **(B)** Pie charts showing individual subset contribution to the whole NK cell population in UCB and APB. Data are from 1 experiment with n=3 UCB and n=3 APB samples.

### Each NK cell population expresses a distinct pattern of activating and inhibitory receptors

We further characterised the identified NK cell populations for their expression of immune checkpoints and NK cell receptors (Figure 5A). CD96 was the highest on CD56^bright^ NK cells (Population A) and on CD56^dim^CD16^-^ KIR^+^ NK cells (Population F_1_) while conventional terminally differentiated NK cells (Population B_2_) and unconventional CD16^-^KIR^-^ terminally differentiated NK cells (Population G_3_) expressed the lowest amounts of CD96. These observations support the hypothesis that CD96 expression decreases with NK cell maturation and might account for the reduced expression of CD96 in APB compared to UCB. Concordant with published data (8), TIGIT expression was low on conventional CD56^bright^CD16^-^ NK cells (Population A_1_). TIGIT expression was the highest on CD56^bright^ CD16^+^ NK cells (Population A_2_) and NKp46^+^KIR^+^CD56^-^ NK cells (Population C_3_). TIM-3 was expressed at very high levels on most NK cell subsets, however low expression was observed on some unconventional subsets (Populations C_2_, C_3_, E_2_, F_2_, G_1_ and G_3_). We observed higher NKG2A expression on CD56^bright^ NK cells (Population A) compared to conventional CD56^dim^ NK cells (Population B), in line with previous reports (6). Interestingly, the intermediate CD56^+^CD16^+^ (Population D) as well as Population H also expressed higher levels of NKG2A compared to other NK cell subsets, possibly indicating a proximity with the CD56^bright^ subsets in terms of progeny. DNAM-1 was expressed on most NK cell subsets with the exception of NKp46^-^PD-1^+^ NK cells (Population E_2_) that were mostly DNAM-1^-^ and Population C_3_ that contained a high proportion of DNAM-1^-^ cells, suggesting a potential progeny link between these two subsets. NKp30 was also expressed on most NK cell subsets except for CD56^neg^ subsets C_2_ and C_3_ and mature NKp46^-^PD-1^+^ subset E_2_. Population E_2_ also displayed low NKG2D expression compared to all the other subsets. In summary, each NK cell subset identified by unsupervised means expressed displayed unique repertoire of receptor expression. We did not observe clear co-expression patterns, indicating receptors expression and segregation between subsets might be independently regulated. Interestingly, NKG2A was the only receptor that was consistently reduced on all subsets in UCB compared to APB, suggesting that low NKG2A expression is a specific feature of UCB NK cells (Supplementary Figure 5). Taken together, our data support a model for NK cell maturation where CD56, NKG2A, CD96, NKp46 and NKG2D expressions are progressively decreased concomitantly with sequential acquisition of CD16, KIR and CD57 markers (Figure 5B).

**Figure 5.**
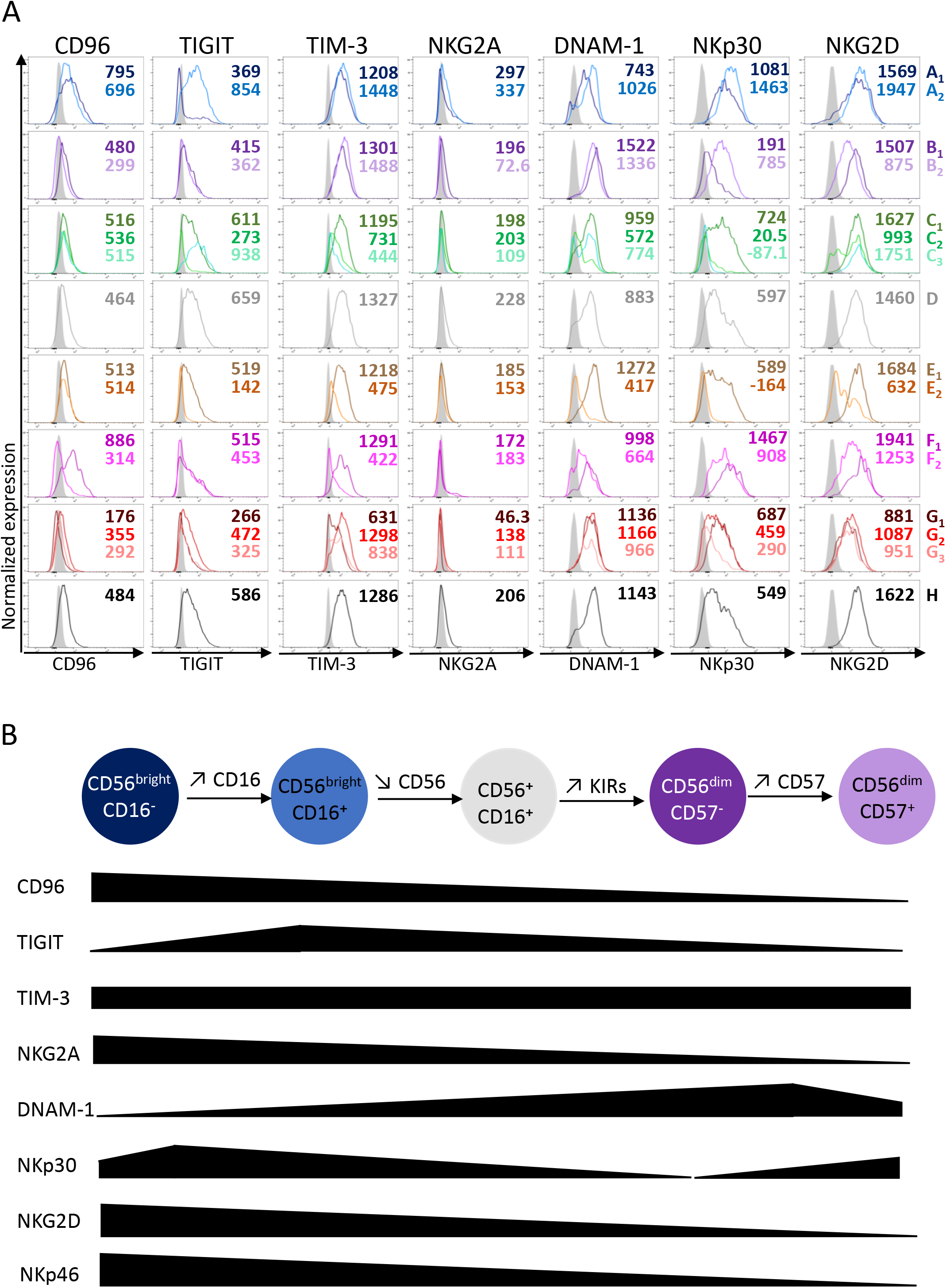
Receptor expression analysis on different NK cell subsets support a linear maturation model from CD56^bright^ to CD56^dim^ NK cells. **(A)** Histograms show NK cell receptor expression on NK cell populations defined as in Figure 3. Numbers indicate geometric mean fluorescence intensity. Data are from 1 experiment with n=3 UCB and n=3 APB pooled samples. **(B)** Proposed linear maturation model showing changes in NK cell receptor expression from CD56^bright^ to CD56^dim^ NK cells.

### Healthy donor NK cells upregulate LAG-3 and PD-1 upon cytokine stimulation

T cells are known to up-regulate immune checkpoints upon activation (26). To determine if immune checkpoints are also upregulated on activated NK cells, we stimulated UCB and APB mononuclear cells with IL-12, IL-15 and IL-18. Cells cultured with IL-15 only to ensure NK cell survival were used as control. The prevalence of TIM-3^+^ cells was modestly, but significantly, up-regulated in stimulated UCB NK cells, while no significant difference was observed in APB (Figure 6A). Interestingly, the frequencies of both TIGIT^+^ and CD96^+^ cells decreased upon IL-12/15/18 stimulation on both UCB and APB NK cells compared to IL-15 only cultures (Figures 6B and 6C), although we noticed that cultured NK cells (IL-15 alone and IL-12/15/18) had up-regulated CD96 expression compared to uncultured NK cells (Supplementary Figure 6A). We observed a slight but significant increase in PD-1+ NK cells in both UCB and APB (Figure 6D). Finally, whilst LAG-3 could not be detected on unstimulated NK cells, LAG-3 expression was consistently induced on a small proportion of NK cells, in both UCB and APB (Figure 6E and 6F). Compared to their LAG-3^-^ counterparts, LAG-3^+^ NK cells in both UCB and APB harboured higher frequencies of TIM-3^+^, PD-1^+^ and CD96^+^ cells, indicating co-expression of these immune checkpoints (Figure 6G). NKG2D was also expressed on a higher percentage of LAG-3^+^ NK cells compared to LAG-3^-^ counterparts. No difference was observed in the frequencies of NKp46^+^ or CD57^+^ cells within the LAG-3^+^ and LAG-3^-^ populations, although small but significant variations in DNAM-1^+^, KIRs^+^ and NKp30^+^ cell frequencies were observed in UCB but not in APB (Supplementary Figure 6B). Collectively, these data indicate that the T cell-associated immune checkpoints PD-1 and LAG-3 are up-regulated on activated NK cells while the nectin-like family receptors TIGIT and CD96 were down-regulated upon cytokine stimulation.

**Figure 6.**
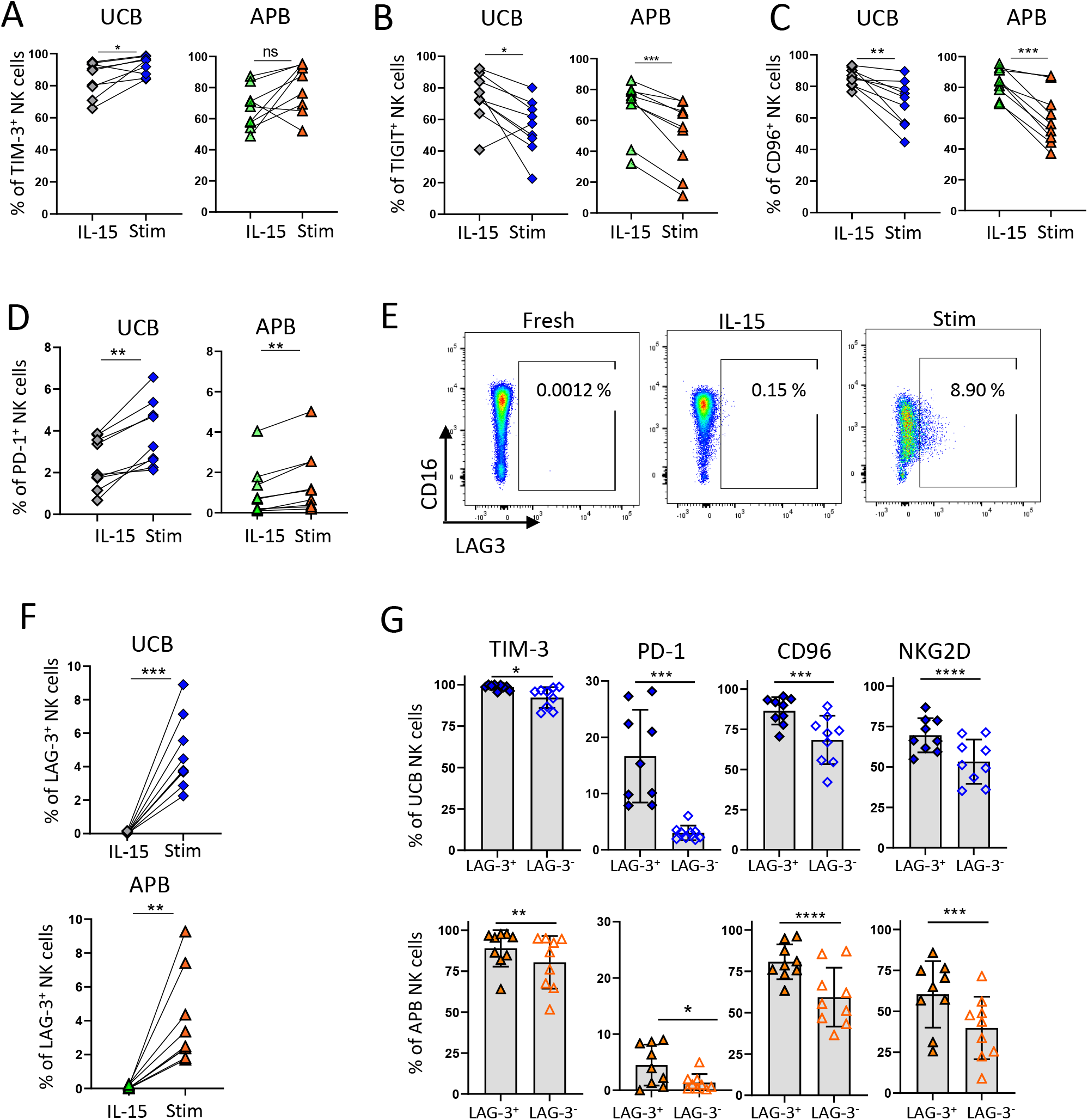
NK cells up-regulate TIM-3, LAG-3 and PD-1 and down-regulate TIGIT and CD96 upon cytokine stimulation. UCB or APB mononuclear cells were cultured for 36 hrs in the presence of 50 ng/mL of IL-15. Cytokine stimulation was performed by the addition 10 ng/mL of IL-12p70 and 10 ng/mL IL-18 after 24 hrs. **(A-D)** Percentages of NK cells positive for TIM-3 (A), TIGIT (B), CD96 (C), or PD-1 (D) in IL-15 only cultures (IL-15) or IL-15, IL-12, IL-15 stimulated NK cells (Stim). **(E)** Representative dot plot showing the percentages of NK cells positive for LAG-3 in uncultured sample (fresh), after 36 hrs culture with IL-15 (IL-15) or after IL-15, IL-12, IL-18 stimulation (Stim). **(F)** Quantification of the data shown in E. **(G)** Percentages of IL-15/12/18 stimulated LAG-3^+^ and LAG-3^-^ NK cells positive for TIM-3, PD-1, CD96 and NKG2D in UCB (top) and APB (bottom). Data are from 3 independent experiments with a total n=9 UCB and n=9 APB samples. Graphs show mean + SD. Data were analysed with a paired t-test. * p<0.05; ** p<0.01; *** p<0.001; **** p <0.0001

### NK cell subsets are conserved in IL-15 cultures and upon cytokine stimulation

Using FlowSOM, we analysed NK cell populations following IL-15 only and IL-12/15/18-stimulation (Figures 7A-D). Despite CD56 up-regulation on cultured NK cells (27), we were able to identify populations equivalent to the main 8 populations defined on unstimulated samples (see supplementary methods). Importantly, population distribution across UCB and APB also matched steady states observations (Figures 4B and 7E). UCB samples accounted for most of the CD56^bright^CD16^+^ NK cells (a_2_), conventional mature CD57^-^ NK cells (b_1_), CD56^neg^ NK cells (c_1_ and c_2_), intermediate CD56^+^CD16^+^ NK cells (d) and mature NKp46^-^ populations (e) while APB samples accounted for most of conventional (b_2_) and unconventional (g) terminally differentiated NK cells (Figure 7E). Overall, these data indicate that differences between UCB and APB in NK cell subsets identified in unstimulated samples are conserved upon short-term IL-15 culture and cytokine stimulation.

**Figure 7.**
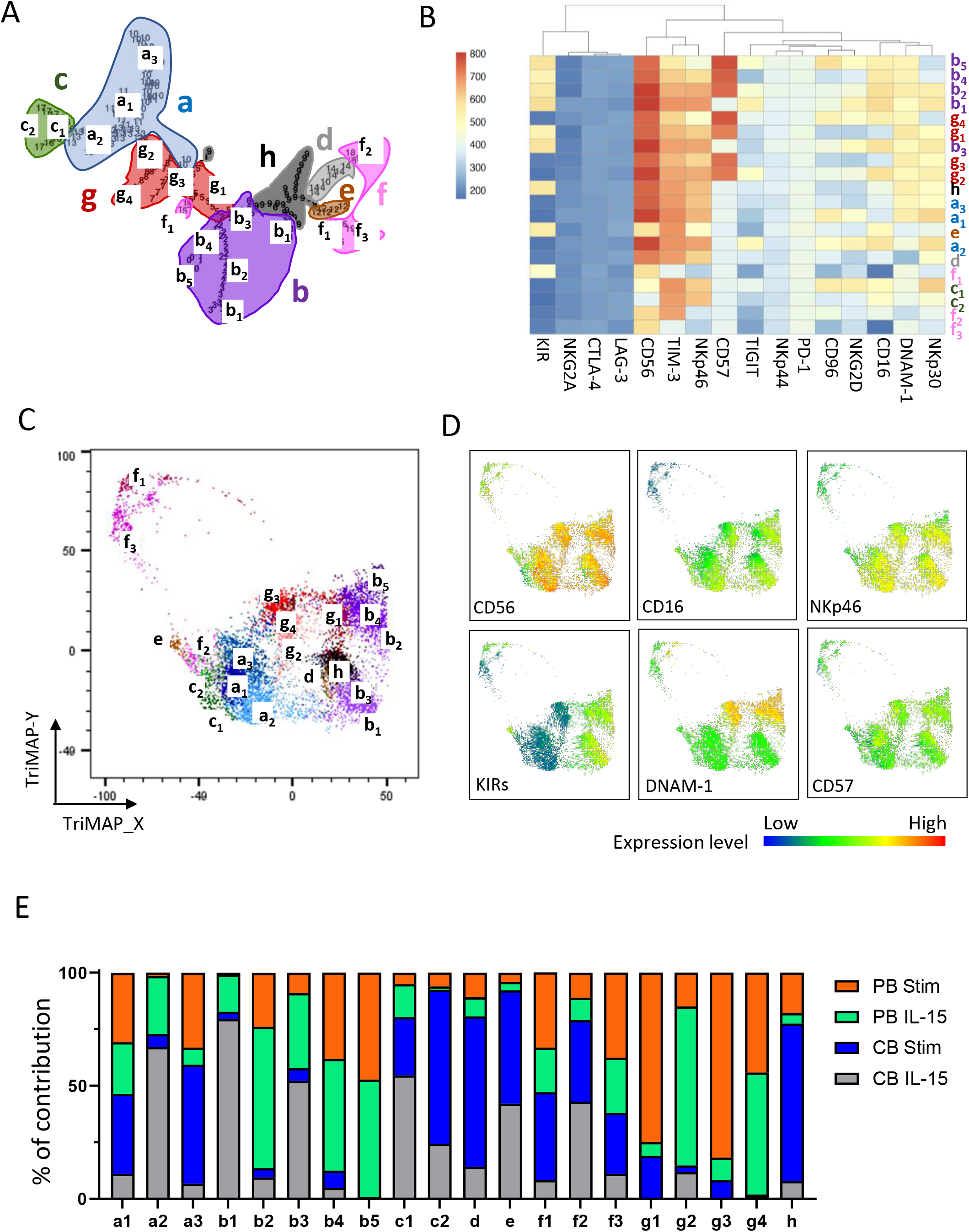
High-dimensional analysis of cytokine-stimulated NK cells identifies the main 8 populations found in uncultured NK cells. FlowSOM was run on NK cells from 3x IL-15 only cultured and 3x IL-12/15/18 stimulated UCB and 3x IL-15 only cultured and 3x IL-12/15/18 stimulated APB samples (1100 events per sample) using a 15×15 matrix and a set metacluster number of 20. Analysis was run on all compensated fluorescent parameters except the viability dye, human CD45, CD3 and CD33/CD19. **(A)** FlowSOM generated tree annotated with the 8 main populations (a-h) corresponding to the proposed stimulated equivalent of populations (A-H) identified in Figure 3. Sub-populations annotated with underscript numbers are shown. **(B)** FlowSOM generated heatmap showing the phenotype of each identified population. **(C)** TriMap algorithm was run on the same dataset and FlowSOM populations were overlaid. **(D)** Heatmaps showing expression levels of NK cell receptors used to identify the different subsets. **(E)** Graph shows the contribution of IL-15 cultured UCB and APB NK cells as well as IL-12/15/18 stimulated UCB and APB NK cells to the identified FlowSOM populations.

### Identification of cytokine-induced changes in receptor expression conserved across multiple NK cell subsets

Analysis of immune checkpoint expression revealed that most NK cell subsets expressed CD96, TIGIT and TIM-3 with the exception of the CD16-negative populations f_1_ and f_3_ (Figure 8A). In addition, TIGIT was also absent on populations e (NKp46^-^ NK cells) and f_2_ (CD16^-^ KIR^-^ NK cells) whilst TIM-3 was absent on the terminally differentiated population b_5_. In line with observations in fresh samples, PD1^+^ NK cells mostly clustered within CD56^neg^ (c_2_) and NKp46^-^ (e) populations. In striking contrast, no cluster of LAG-3 cells was observed, which indicates that LAG-3 is up-regulated sporadically on multiple NK cell subsets. These results suggest that immune checkpoints are not co-regulated in NK cells and different immune checkpoints follow different patterns of expression ranging from pan-expression on the majority of NK cell subsets (TIM-3, TIGIT, CD96), to segregated expression on specific subsets (PD-1) or sporadic up-regulation across multiple subsets (LAG-3). Further comparison between IL-15 cultures and IL-12/15/18-stimulated NK cells revealed that cytokine stimulation resulted in increased frequency of DNAM-1^+^ cells in both UCB and APB NK cells, with decreased frequencies of NKG2D^+^, NKp30^+^ and CD16^+^ cells (Figure 8B). Of note, no differences were observed regarding CD57, KIR or NKp46 expression (Supplementary Figure 7). Analysis of expression on specific populations revealed that (with the exception of TIGIT) a given marker was regulated in the same direction in all NK subsets (i.e. up-regulated on all subsets with activation or down-regulated on all subsets with activation; Figure 8C). No or negligible differences were observed for NKp30, CD96 and TIGIT expression between IL-15 and stimulated NK cell subsets. These results suggests that modification in NKp30, CD96 and TIGIT expression observed at the whole NK cell population level reflects changes in the representation of NK cell subsets. By contrast, CD16 and NKG2D were down-regulated on multiple NK cell subsets in both UCB and APB while DNAM-1 and TIM-3 were up-regulated (Figure 8C). Based on these data, we propose that there exists a common response to cytokine stimulation shared across multiple NK cell subsets and characterised by the up-regulation of LAG-3, TIM-3 and DNAM-1 and the downregulation of NKG2D and CD16.

**Figure 8.**
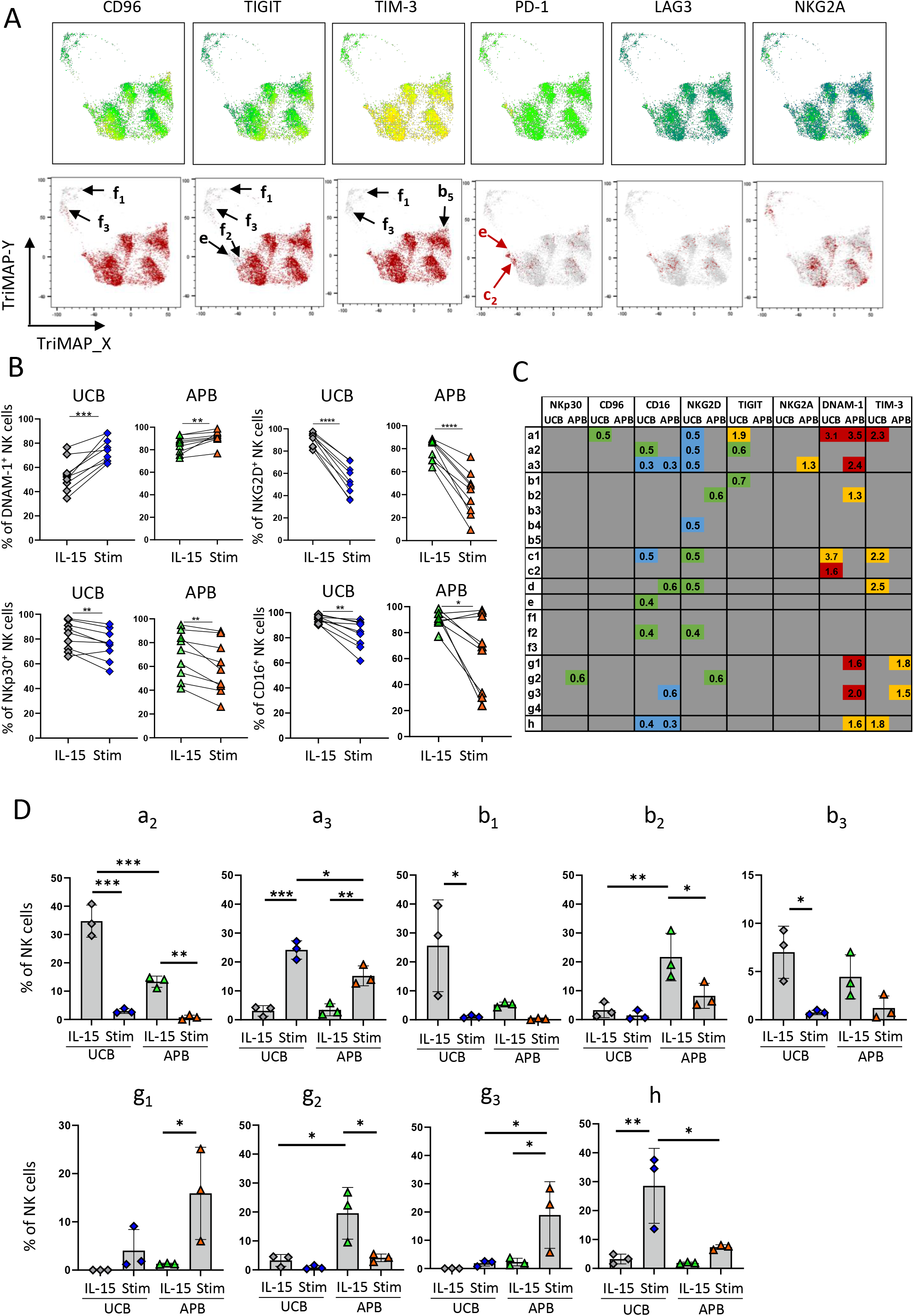
UCB and APB NK cell populations exhibit similar changes in receptor expression upon cytokine stimulation. FlowSOM analysis was run as per Figure 7. **(A)** Heatmaps showing expression levels of inhibitory receptors (top). Cells positive for a given receptor (when gating on the positive population based on control staining) are shown in red (bottom). Black arrows point to populations negative for CD96, TIGIT or TIM-3 while red arrows indicate populations positive for PD-1. **(B)** Percentages of NK cells positive for DNAM-1, CD16, NKG2D and NKp30 in IL-15 only cultures (IL-15) or IL-12/15/18 stimulated NK cells (Stim). **(C)** Heatmap showing variations in geometric mean fluorescence intensity for a given marker between IL-12/15/18 stimulated and IL-15 cultured NK cells. Numbers indicate fold increase. Markers increased in stimulated populations with p<0.05 or p<0.01 are shown in yellow and red, respectively. Markers decreased in stimulated populations with p<0.05 or p<0.01 are shown in green and blue, respectively. **(D)** Contribution of specific NK cell populations to the whole NK cell population in UCB and APB IL-15 only cultures (IL-15) or IL-12/15/18 stimulated NK cells (Stim). Graphs show mean + SD. n=3 samples per group. Data were analysed with a paired t-test (B), multiple t-tests (C) or one-way Anova followed by Sidak’s multiple comparison test (D). * p<0.05; ** p<0.01; *** p<0.001; **** p <0.0001

### Differentiation of distinct CD16^+^TIGIT^+^ NK cells subsets into corresponding CD16^-^TIGIT^-^ NK cells subsets upon cytokine stimulation

Finally, we analysed the effect of cytokine stimulation on subset distribution (Figures 8D). In both UCB and APB cultures, IL-12/15/18 stimulation led to a reduction in CD56^bright^CD16^+^ NK cells (a_2_ subset) concomitant to an increased representation of a new CD56^bright^CD16^-^ subset (a_3_) that differed from the steady-state identified CD56^bright^CD16^-^ subset (A_1_/a_1_) by its low expression of CD96 and NKG2D. Conventional mature NK cells (b_1_ and b_3_ in UCB and b_2_ in APB) were also reduced upon stimulation, as were the unconventional CD16^+^KIR^-^ terminally differentiated NK cells (g_2_). By contrast, representation of unconventional CD56^bright^KIR^+^ NK cells that were negative for CD16 (h) was increased upon stimulation of UCB. Similarly, representation of CD16^-^ unconventional populations (g_1_ and g_3_) was increased upon cytokine stimulation in APB. Interestingly, cytokine stimulation was associated with a decreased representation of TIGIT^+^ subsets (a_2_, b_1_, b_2_, g_3_) and increased representation of TIGIT^-^ subsets (a_3_, h, g_1_, g_2_). Subsets increased over activation also arbored reduced levels of CD96 (a_3_ compared to a_2_; h compared to b_1_/b_3_; g_1_/g_3_ compared to b_2_/g_2_) as well as NKp30 expression (a_3_ compared to a_2_ and g_1_/g_3_ compared to b_2_/g_2_). Overall, these data support a model where distinct CD16^+^TIGIT^+^ NK cell subsets characterised by differential expression of CD56, KIRs and CD57 differentiate into new CD16^-^ TIGIT^-^ NK cell subsets with reduced CD96 and NKp30 expression upon cytokine stimulation (Supplementary Figure 8).

## Discussion

In this study, we provide the first in depth characterisation of NK cell populations in UCB, an attractive source for cellular therapy (15). Through high-dimensional flow cytometric analysis combined with unsupervised clustering analysis, we identified distinct NK cell populations in UBC and APB and showed that these populations were preserved upon stimulation with IL-12/15/18, a protocol that has been used to generate memory NK cells with increased clinical potential (28). Herein, we have also identified conserved phenotypic alterations in response to IL-12/15/18 stimulation that were shared across different NK cell subsets in both UCB and APB.

The regulation NK cell functions through immune checkpoints has recently generated much interest given its importance to clinical outcomes (24). Our data indicate that TIM-3, TIGIT and CD96 are the main immune checkpoints expressed by unstimulated healthy donor NK cells, an observation that corroborates previous reports (22, 24, 29). Interestingly, while immune checkpoints such as TIGIT are up-regulated on activated or exhausted T cells (30), we found that TIGIT and CD96 were down-regulated upon cytokine stimulation of NK cells. This suggest that TIGIT and CD96 might play different roles in T cells and NK cells. Two other immune checkpoints, LAG-3 and PD-1, were up-regulated post-stimulation, supporting a positive association of these markers with NK cell exhaustion (21). No LAG-3 expression was observed in the steady state and LAG-3^+^ NK cells identified post-stimulation displayed enriched expression of PD-1 and importantly were detected across multiple NK cell subsets. By contrast, PD-1 was also found on unstimulated UCB NK cells, with the expression being restricted mainly to CD56^neg^ and mature NKp46^-^ subsets. Whether PD-1 might regulate these UCB-specific populations and its implications on therapeutic use of UCB NK cells warrants further investigation. Collectively, our study reveals enrichment of immune checkpoints TIGIT, CD96, TIM-3 and PD-1 on UCB NK cells, possibly accounting for the decreased reactivity of unstimulated UCB NK cells compared to their APB counterparts (17).

In agreement to earlier studies (31, 32), we observed a population of CD56^neg^ NK cells in UCB. These cells have been shown to exhibit limited lytic activity and antiviral functions (31, 32). Here, we provide an exhaustive phenotypic characterization of CD56^neg^ UCB, establishing that as a population they have reduced frequencies of cells expressing DNAM-1, NKG2D, NKp30 and TIGIT compared to the whole UCB NK cell population. CD56^neg^ UCB NK cells have been shown to differentiate into CD56^+^CD16^+^ cells in vitro (32). Surprisingly, we found that CD56^neg^ NK cells lack NKG2A expression, a finding that argues against an immature status of these cells. Alternatively, this finding might indicate that CD56^neg^ NK cells belong to lineage separate to that of CD56^bright^ NK cells, with both CD56^neg^ and CD56^bright^ NK cells possessing the capacity to differentiate into CD56^dim^ NK cells. Our high dimensional analysis revealed further diversity of the CD56^neg^ UCB NK cell population, with differential NKp46, KIRs and PD-1 expression. Via unsupervised clustering analysis, we also identified another population of mature NK cells characterised by a CD16^+^KIR^+^NKp46^-^ phenotype that was mostly found in UCB. These NKp46^-^ mature NK cells were also CD56^low^ and shared many features with the CD56^neg^ NK cells including low expression of DNAM-1 and NKp30 and enrichment in PD-1^+^ cells. Based on these data, we hypothesise that CD56^neg^ and NKp46^-^ mature NK cells are related. Our results also suggest that these cells are not any intermediate in the linear maturation model of NK cells where CD56^bright^ differentiate into CD56^dim^ NK cells (6). CD56^neg^ and NKp46^-^ mature NK cells might represent a branched differentiation pathway (Supplementary Figure 8) or even belong to a distinct lineage.

Our analysis of IL-15 cultured and IL-12/15/18 activated NK cells revealed that the identified NK cell subsets in UCB and APB were maintained upon activation. Populations could notably be identified using the maturation markers NKG2A, KIRs and CD57 that appeared stable in culture. These data indicate that cytokine stimulation did not accelerate NK cell maturation. Instead, cytokine stimulation induced a loss of CD16 expression in multiple NK cell subsets leading to the appearance of corresponding CD16^-^ subsets. CD16 shedding on activated NK cells has been described and is mediated by A disintegrin and metalloprotease-17 (ADAM17) (33). To our knowledge, this study is the first one to demonstrate that CD16 loss upon activation is a phenomenon shared by distinct NK cell subsets including CD56^bright^CD16^+^ and CD56^+^CD16^+^ that were more represented in UCB. Moreover, we show that CD16 loss is concomitant to the loss of TIGIT on NK cell surface and associated with decreased NKp30 and CD96 as well as increased DNAM-1 expression.

Our results have uncovered major differences between UCB and APB NK cells including distinct NK cell populations and higher expression of immune checkpoints in UCB NK cells. These differences need to be considered in the choice of NK cell source for adoptive therapy.

## Acknowledgements

This work was supported by the Mater Foundation, Brisbane, Australia and by an Innovator Grant from the Queensland Children Hospital Foundation attributed to CG (RPC00037). CG is supported by a Discovery Early Career Research Award (DECRA) from the Australian Research Council (ARC) (DE210101144).

## Contribution

Study conception and design: CG; Experimental design: IB, VW, DPS and CG; Data acquisition: IB; Data analysis: IB and CG; Data interpretation: IB, DPS, KR and CG; Figure conception and design: IB and CG; Manuscript writing: IB and CG; Manuscript editing: IB, DPS, KR and CG. All authors read and approved the final manuscript.

