## Supplementary data for "High dimensional flow cytometric analysis reveals distinct NK cell subsets but conserved response to cytokine stimulation in umbilical cord blood and adult peripheral blood"

### Supplementary methods.

**Cell isolation**. Cell isolation from adult peripheral blood (APB) and umbilical cord blood (UBC) samples was performed in sterile conditions at room temperature. Firstly, blood was diluted with equal amount of PBS to a total of 35ml and underlayed with 15ml of blood cell density gradient medium Ficoll-Paque Plus (GE Healthcare Pharmacia, 17-1440-02) in a 50ml conical tube. Cells were centrifuged at 1600 RPM for 20 minutes without break to prevent disruption of the density gradient. The mononuclear cells from plasma/Ficoll interface were collected using a sterile transfer pipette and washed twice with PBS.

**Cryopreservation.** PBMCs were cryopreserved in 90% FBS and 10% DMSO solution. Cord blood MNCs were cryopreserved using 90% plasma and 10% DMSO solutions. Cryovials were placed in NALGENE Mr.Frosty in -80° C for 12-24 hours and subsequently relocated into long term -196° C liquid nitrogen storage. Cells were frozen at maximum concentration of $50x10^6$ cells/ml in 1ml cryovials.

**Thawing.** Cryopreserved cells were rapidly thawed at 37° C and washed once in complete RPMI-1640 medium (Invitrogen, Grand Island, NY) supplemented with 10% heat inactivated Fetal Calf Serum (FBS) (Gibco cat #, 1099141), 2 mM l-glutamine (Gibco–Invitrogen, Grand Island, NY), 100 µg/mL Penicillin-100 U/mL streptomycin (Sigma, St. Louis, MO), and 10 mM HEPES (Sigma) and RNase free DNase I (final concentration of 12.5 µg/ml, Roche Diagnostics, Mannheim, Germany). Cells were centrifuged at 1600 rpm, 5 minutes, 20° C. Following this, cells were resuspended in complete RPMI-1640 medium containing RNase free DNase I (final concentration of 6.25 µg/ml) and incubated for 30 minutes at 37° C. Cells were filtered using 70-micron sterile filter, washed, and resuspended in a complete medium.

**Cell culture.** Mononuclear cells were cultured at $1x10^6$ cells/ml for 36 hrs in Corning cell culture flasks at 37°C, 5% CO2 in complete RPMI-1640 medium (Gibco–Invitrogen) supplemented with 10% heat inactivated Fetal Calf Serum (FBS) (Gibco–Invitrogen cat #, 1099141), 2 mM L-glutamine (Gibco–Invitrogen), 100 µg/mL Penicillin-100 U/mL streptomycin (Sigma-Aldrich), 10 mM HEPES (Sigma-Aldrich), $1x10^{-4}$M non-essential amino acids (Gibco–Invitrogen) and 1mM of sodium pyruvate (Gibco–Invitrogen). For cell survival with 50 ng/ml animal-free recombinant human IL-15 (Peprotech) was added. For stimulation, 10ng/ml, human IL-12p70 (HEK293) (Peprotech) and 50ng/ml recombinant human IL-18 (MBL International) were added to the IL-15 cultures for the last 12 hrs (overnight stimulation).

**Hierarchical clustering analyses**. Data was analysed using Flowjo v10.7.2. NK cells were gated as per Supplementary Figure 1A and NK cells from samples to be analysed were concatenated. Same numbers of NK cells per sample were exported for the analysis. 10,000 events per sample were exported for data shown in Figure 3 and Supplementary Figures 3A-B; 7,000 events per sample were exported for data shown in Supplementary Figure 3C and 1,100 events per sample were exported for data shown in Figure 7 and Figure 8. FlowSOM v2.2.0 using R v4.1.2. was run on the concatenated sample. Analysis was run on all compensated fluorescent parameters except the viability dye, CD3 and CD33/CD19. Human CD45 was also excluded from the analysis for data presented in Supplementary Figure 3C, Figure 7 and Figure 8. The number of metaclusters was set to 20, using a 10x10 matrix (data presented in Figure 3 and Supplementary Figure 3) or a 15x15 matrix (data presented in Figure 7 and Figure 8; as the 10x10 matrix failed to generate 20 metaclusters).

**Population identification**. NK cell populations were determined using FlowSOM generated heatmap and tree, regrouping together populations with similar phenotype and, when possible regrouping populations that appeared close together on the tree. Rationale for the identification of Figure 7 populations corresponding to the cultured equivalents of Figure 3 populations was as followed. Considering that CD56 is up-regulated upon culture, $CD56^{bright}$

NK cells (Population a) was further defined by the absence of KIR and CD57 expression while conventional $CD56^{dim}$ NK cells (Population b) would have become $CD56^{+/bright}$ and could be identified through their expression of KIR and CD16. $CD56^{neg}$ NK cells (Population c) were DNAM-$1^{low}$, a feature reminiscent of the unstimulated Population $C_3$. Population d was defined as $CD56^{+}CD16^{low}$ which, similarly to unstimulated population D was $KIR^{-}$. We identified a population of mature $NKp46^{-/low}KIR^{+}$ NK cells (population e) that appeared similar to unstimulated population E. Similarly to unstimulated Population F, unconventional $CD56^{dim}$ NK cells (Population f) were $CD16^{-}CD57^{-}$ and were comprised of $KIR^{+}$ and $KIR^{-}$ populations. We also observed unconventional terminally differentiated NK cells (Population g) that were $CD57^{+}$ and, like unstimulated population G could be divided into $CD16^{-}KIR^{+}$, $CD16^{+}KIR^{-}$ or $CD16^{-}KIR^{-}$ cells. Finally, we identified an equivalent to unstimulated population H ($CD56^{bright}$ $KIR^{-}$) that were further defined by their low expression of CD16 (Population h).

**Mean Fluorescence Intensity (MFI) fold increase analysis**. For Figure 8C, geometric MFI of FlowSOM generated populations were exported after gating on individual samples. Populations containing less than 10 events were excluded. Data was compiled on Excel and MFI for IL-12/15/18 stimulated versus IL-15 cultured NK cells were compared on prism using multiple t-test.

**Supplementary Table1. Antibodies used.**

| Specificity | Conjugate /dye | Channel | Clone | Optimal dilution | Company | Catalogue number |
|---|---|---|---|---|---|---|
| LAG-3 | BUV395 | UV_379/28 | T47-530 | 1/50 | BD | 745640 |
| CD16 | BUV496 | UV_515/30 | 3G8 | 1/200 | BD | 612944 |
| DNAM-1 | BUV615 | UV_610/20 | DX11 | 1/25 | BD | 751233 |
| NKp30 | BUV661 | UV_670/30 | P30-15 | 1/100 | BD | 750510 |
| NKG2D | BUV737 | UV_740/35 | 1D11 | 1/100 | BD | 748426 |
| huCD45 | BUV805 | UV_820/60 | HI30 | 1/200 | BD | 612891 |
| TIGIT | BV421 | V_450/50 | 741182 | 1/400 | BD | 747844 |
| NKG2A | BV480 | V_525/50 | 131411 | 1/50 | BD | 747923 |
| Dead cells | LiveDead FVS 575V | V_610/20 | N/A | 1/500 | BD | 565694 |
| TIM-3 | BV786 | V_780/60 | 7D3 | 1/100 | BD | 742857 |
| CD96 | BB515 | B_515/20 | 6F9 | 1/50 | BD | 564774 |
| NKp44 | BB660-P2 | B_670/30 | P44-8 | 1/100 | BD | 624295 |
| CTLA-4 | BB700 | B_710/50 | BNI3 |  | BD | 566901 |
| CD56 | BB790-P2 | B_780/60 | NCAM16.2 | 1/1000 | BD | 624296 |
| CD158a (KIR2DL1) | PE | YG_586/15 | HP-3E3 | 1/100 | BD | 556063 |
| CD158b (KIR2DL2/3) | PE | YG_586/15 | CH-L | 1/100 | BD | 559785 |
| NKB1 (KIR3DL1) | PE | YG_586/15 | DX9 | 1/200 | BD | 555967 |
| PD-1 | PE-CF594 | YG_610/20 | EH12.1 | 1/100 | BD | 565024 |
| CD33 | PE-Cy5 | YG_670/30 | WM53 | 1/200 | BD | 551377 |
| CD19 | PE-Cy5 | YG_670/30 | HIB19 | 1/200 | BD | 555414 |
| NKp46 | PE-Cy7 | YG_780/60 | 9E2 | 1/100 | BD | 562101 |
| CD57 | APC | R_670/30 | NK-1 | 1/5000 | BD | 560845 |
| CD3 | APC-H7 | R_780/60 | SK7 | 1/200 | BD | 560176 |

### Supplementary Figure Legends

**Supplementary Figure 1. (A)** UCB MNCs were stained with a 20-colour flow cytometry panel. After excluding doublets and gating on human $CD45^+$ live cells, NK cells were defined as $CD3^-CD33^-CD19^-$ cells that were either positive for CD16 or CD56. A representative UCB sample is shown. **(B-C)** UCB and APB MNCs were stained as in (A). (B) Graphs show NK cell percentages from live human $CD45^+$ cells. (C) Graphs show percentages of NK cell positive for a given marker in UCB and APB. **(D)** Percentages of PD-$1^+$ and PD-$1^-$ UCB NK cells positive for a given marker. Data were pooled from 3 independent experiments with a total n=10 UCB and n=9 APB samples and presented as mean $\pm$ SD. Data were analysed with a paired (D) or unpaired (B-C) t-test. ns = non-significant; * p<0.05; ** p<0.01; *** p<0.001; **** p <0.0001

**Supplementary Figure 2. (A)** Percentages of $CD56^{bright}CD16^-$, $CD56^+CD16^+$ and $CD56^{dim}CD16^-$ NK cell subsets out of live human $CD45^+$ cells in UCB and APB. **(B)** Percentages of UCB $CD56^{neg}$ NK cells positive for a given marker compared to the whole UCB NK cell population. Data are from 3 independent experiments with a total n=10 UCB and n=9 APB samples. Graphs show mean $\pm$ SD. Data were analysed with a paired (B) or unpaired (A) t-test. ns = non-significant.

**Supplementary Figure 3. (A)** FlowSOM algorithm was run on NK cells from 3 UCB and 3 APB samples (10,000 events per sample) using a 10x10 matrix and a set metacluster number of 20. Analysis was run on all compensated fluorescent parameters except the viability dye, CD3 and CD33/CD19. FlowSOM generated heatmap is shown. FlowSOM defined populations (numbers) and regrouped populations as defined in Figure 3 (letters) are shown on the right. Populations 5 and 10 were grouped as $CD16^-KIR^-CD57^+$ NK cells ($G_3$), populations 6 and 7 were grouped as $CD16^+KIR^-CD57^+$ NK cells ($G_2$), populations 13 and 15 were grouped as

CD56$^{neg}$NKp46$^{+}$KIR$^{-}$ NK cells ($C_1$) and populations 18 and 19 were grouped as CD56$^{bright}$CD16$^{+}$ NK cells ($A_2$). **(B)** FlowSOM generated tree corresponding to the analysis in (A). The 8 main NK cell populations identified are overlayed. Populations A, D and C at the top of the tree can be distinguished by decreased expression of CD56 (from bright to negative). Clusters of KIR$^{+}$ NK cells and CD57$^{+}$ are identified by black boxes with plain and dashed contour lines, respectively. **(C)** FlowSOM algorithm was run on another set of samples comprising 4 UCB and 4 APB. 7000 NK cells per sample were included in the analysis, using a 10x10 matrix and a set metacluster number of 20. Analysis was run on all compensated fluorescent parameters except the viability dye, CD3, CD33/CD19 and human CD45. Populations with similar phenotype to those previously identified in A were overlaid on the FlowSOM generated tree.

**Supplementary Figure 4.** Pie charts showing individual subset contribution to the whole NK cell population in UCB and APB using the dataset analysed in Supplementary Figure 3C. Data are from 1 experiment with n=4 UCB and n=4 APB samples.

**Supplementary Figure 5.** Graphs comparing the geometric mean fluorescent intensity (MFI) of NK cell receptors between umbilical cord blood (circles, grey bars) and adult peripheral blood (squares, red bars) on NK cell populations. FlowSOM populations were defined as in Figure 3. Data are shown as mean $\pm$ SEM of n=3 UCB and n=3 APB samples.

**Supplementary Figure 6.** UCB or APB mononuclear cells were cultured for 36 hrs in the presence of 50 ng/mL of IL-15. Cytokine stimulation was performed by the addition 10 ng/mL of IL-12p70 and 10 ng/mL IL-18 after 24 hrs. **(A)** Histogram showing CD96 expression on uncultured freshly thawed (black), IL-15 cultured (blue) or IL-15/12/18 stimulated (orange) UCB NK cells. Filled histogram shows control staining with NK cell-defining markers only. Numbers indicate geometric mean fluorescence intensity values. **(B)** Percentages of IL-15/12/18 stimulated LAG-3$^{+}$ and LAG-3$^{-}$ NK cells positive for TIM-3, PD-1, CD96 and NKG2D in UCB (top) and APB (bottom). Data are from 3 independent experiments with a total n=9 UCB and n=9 APB samples. Graphs show mean $\pm$ SD. Data were analysed with a paired t-test. * p<0.05; ** p<0.01; *** p<0.001; **** p <0.0001

**Supplementary Figure 7.** UCB or APB mononuclear cells were cultured for 36 hrs in the presence of 50 ng/mL of IL-15. Cytokine stimulation was performed by the addition 10 ng/mL of IL-12p70 and 10 ng/mL IL-18 after 24 hrs. Percentages of NK cells positive for CD57, KIRs and NKp46 in IL-15 only cultures (IL-15) or IL-15, IL-12, IL-15 stimulated NK cells (Stim). Graphs show mean $\pm$ SD of n=9 samples per group. Data were analysed with a paired t-test. ns= non significant.

**Supplementary Figure 8.** Proposed relationship between NK cell populations identified in Figures 3 and 7 are shown. Pathways are hypothesised based on current literature knowledge (e.g. $CD56^{bright}$ differentiating in $CD56^{dim}$ NK cells and UCB $CD56^{neg}$ NK cells acquiring CD56 expression in culture) and data generated through the present study. Plain arrows show previously demonstrated differentiation pathways and dotted arrows indicate hypothesised differentiation pathways. Black arrows indicate differentiation taken place at steady state, green arrows indicate differentiation promoted through IL-15 culture and red arrows indicate differentiation promoted through cytokine stimulation with IL-12, IL-15 and IL-18.
