## Supplementary figures and images for "High dimensional flow cytometric analysis reveals distinct NK cell subsets but conserved response to cytokine stimulation in umbilical cord blood and adult peripheral blood"

### Supplementary Figures

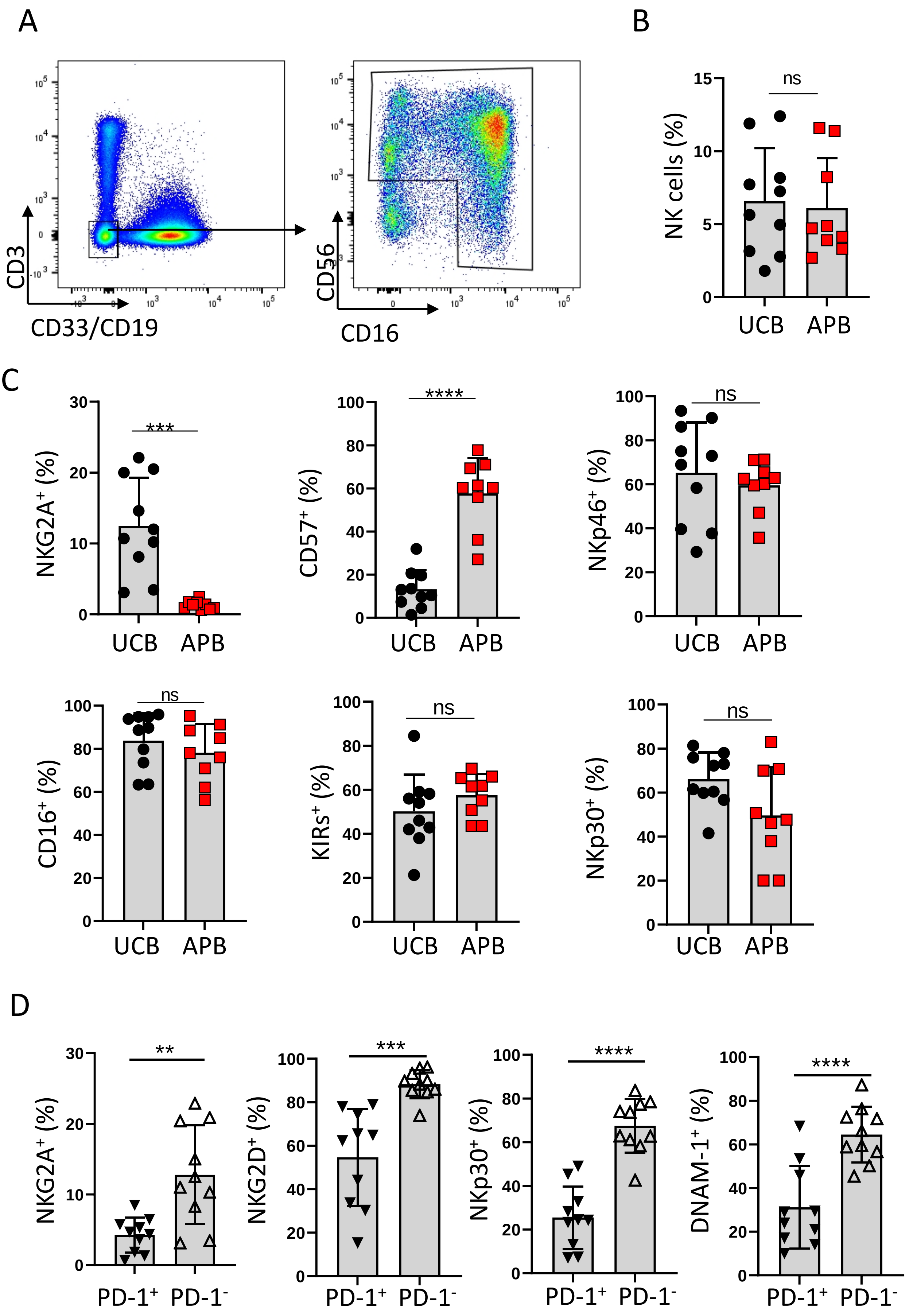

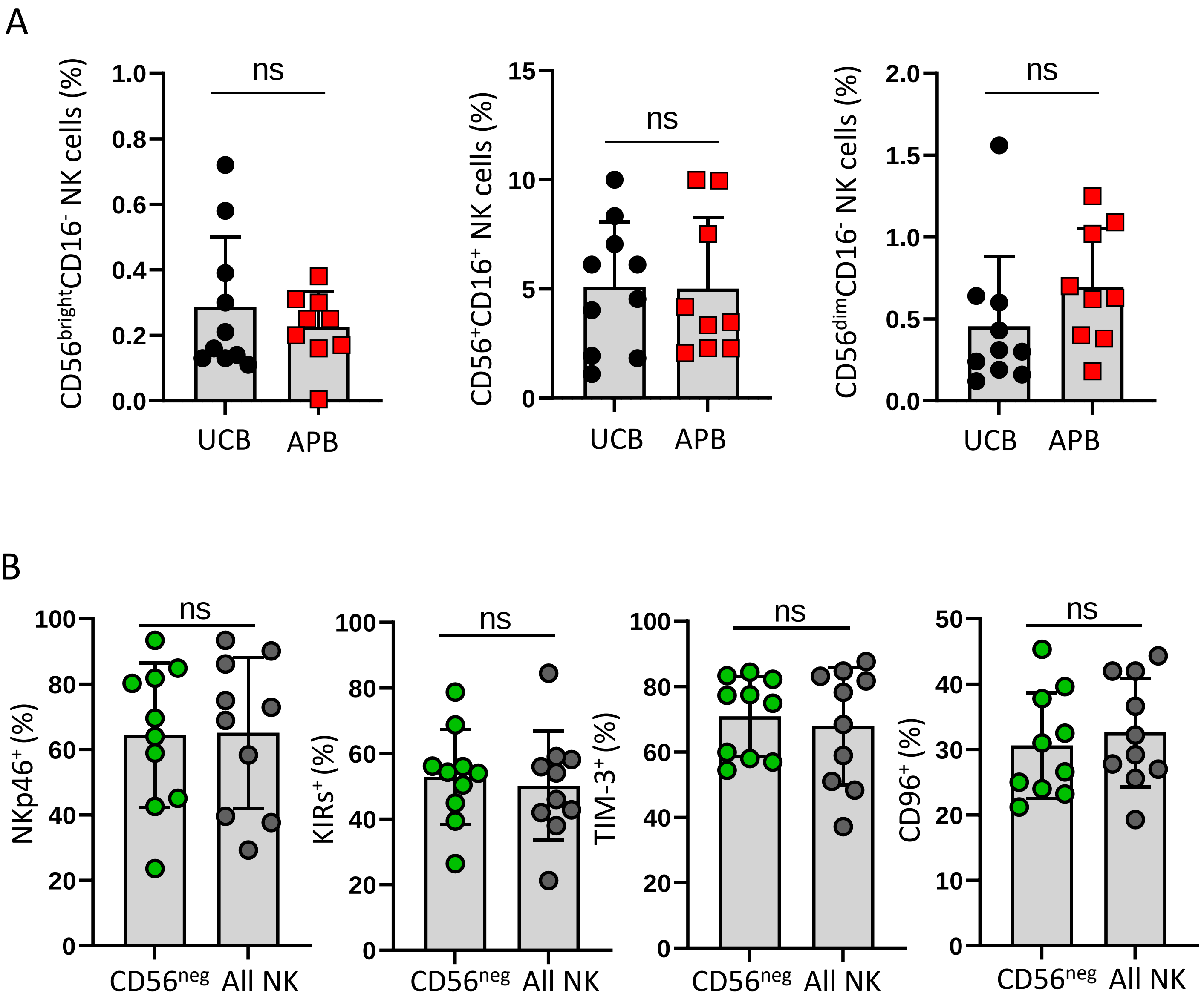

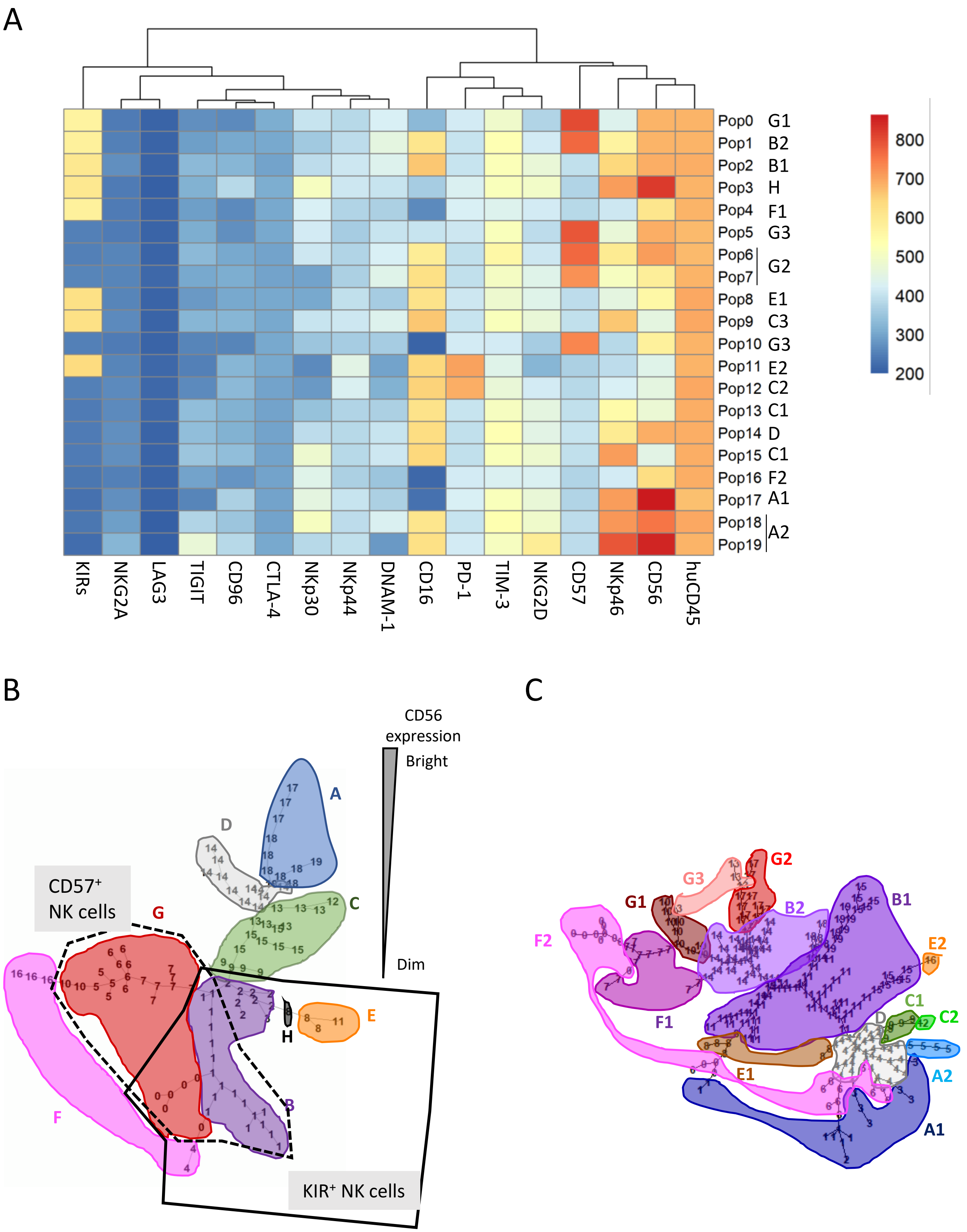

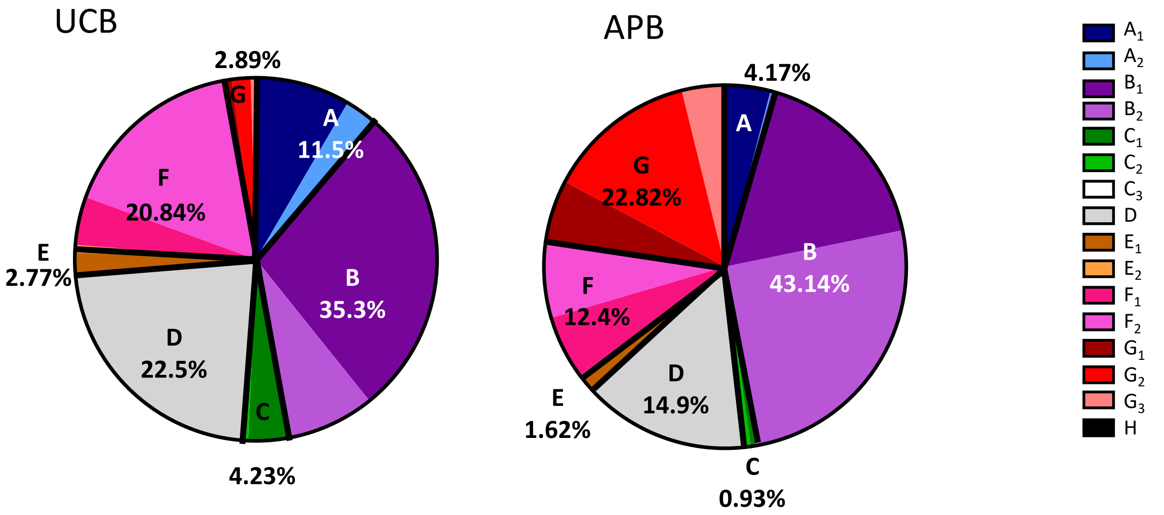

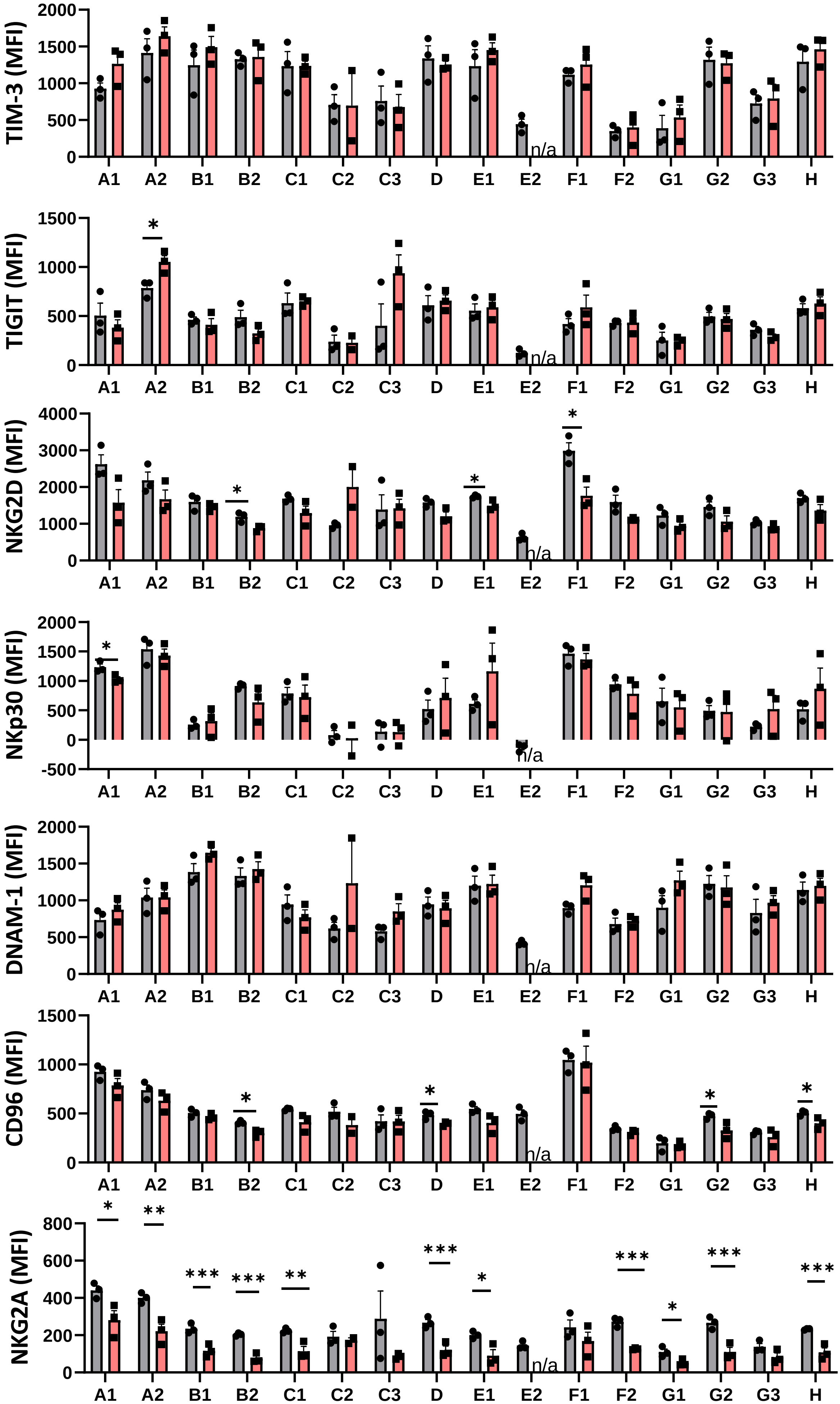

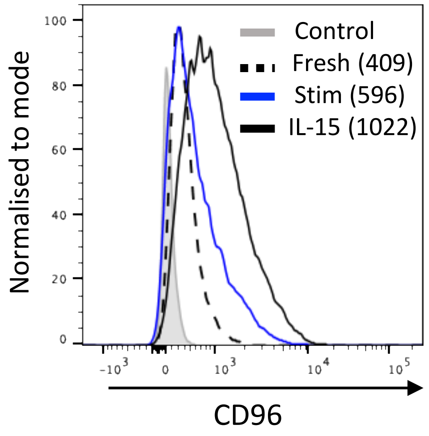

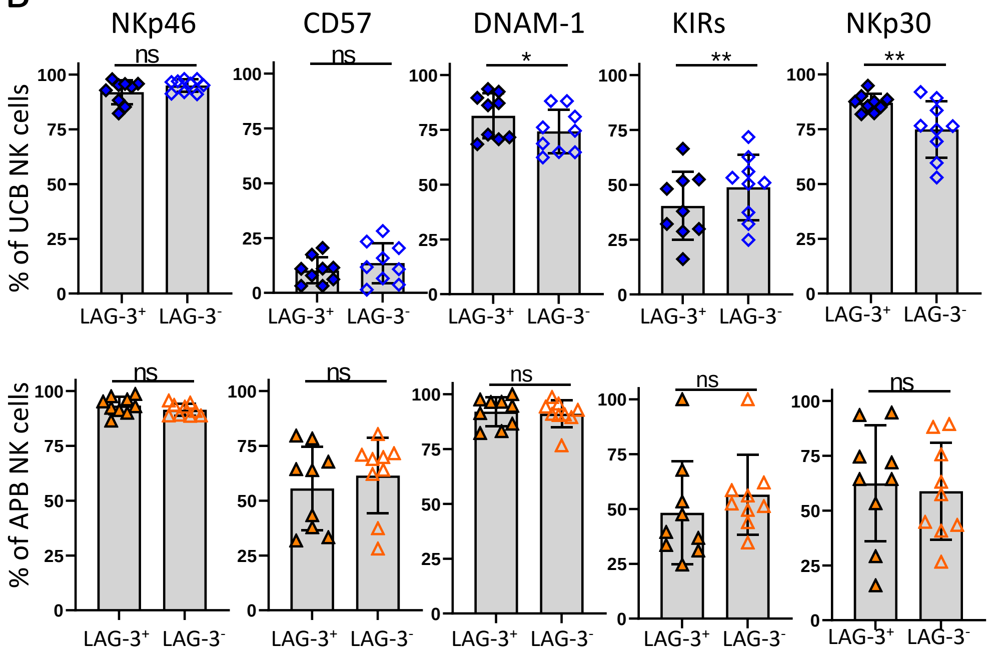

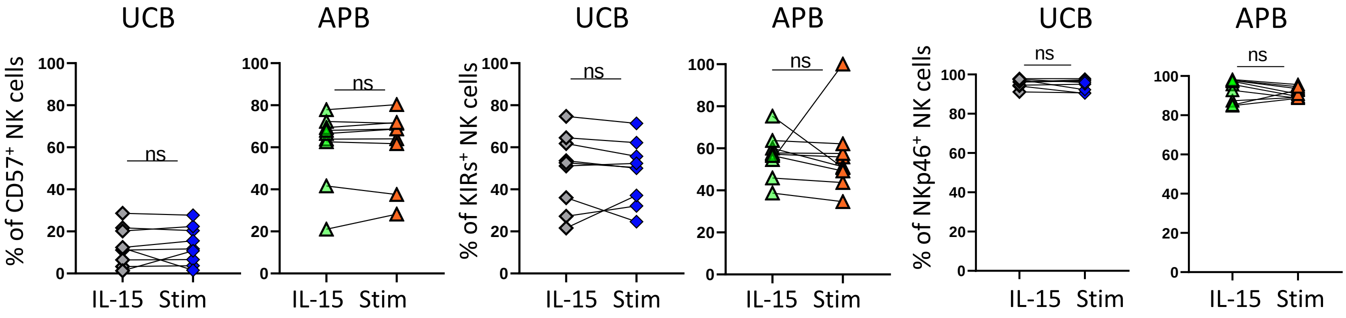

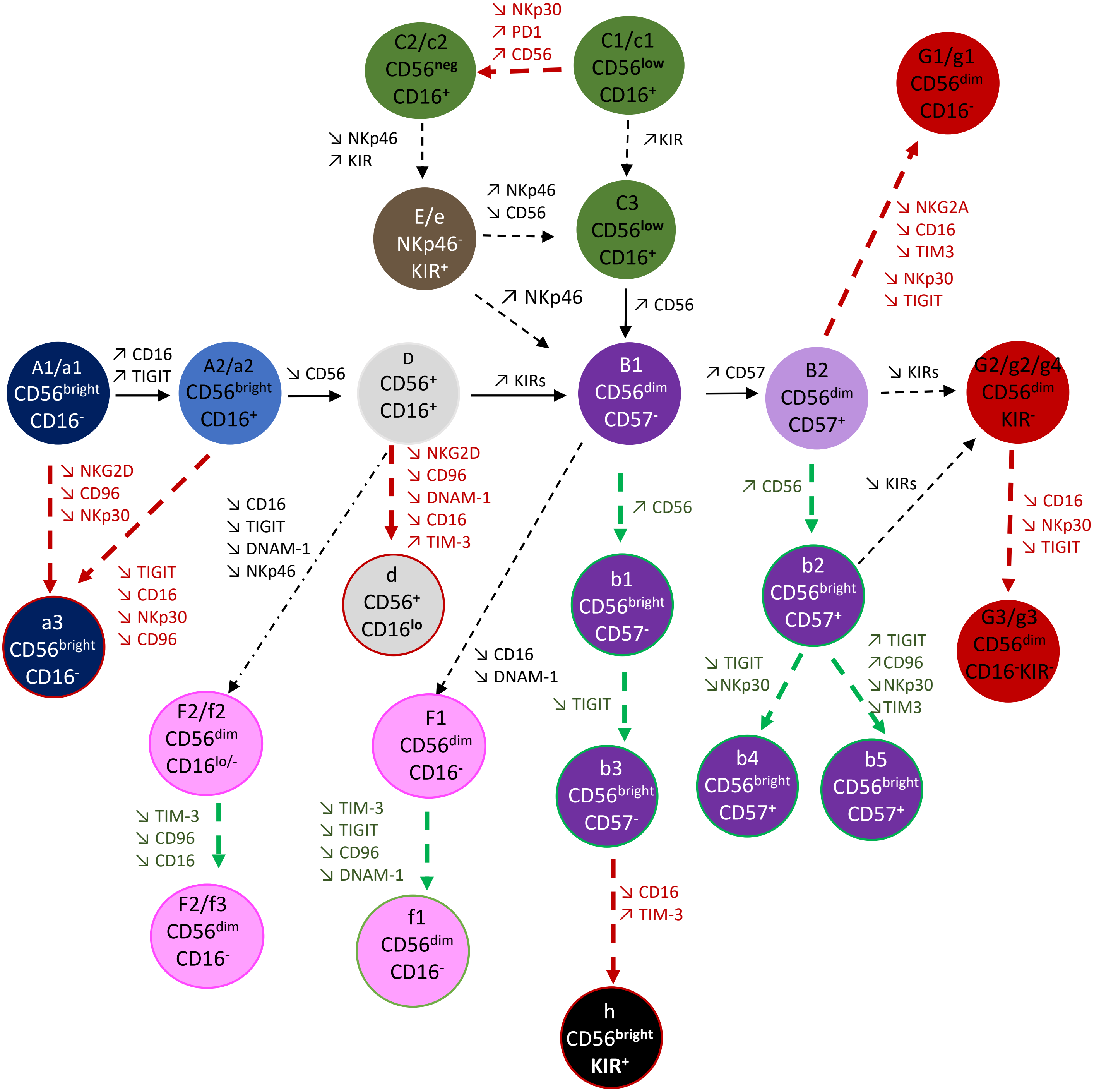
